# Cultivation and genomic analysis of *Candidatus* Nitrosocaldus islandicus, a novel obligately thermophilic ammonia-oxidizing *Thaumarchaeon*

**DOI:** 10.1101/235028

**Authors:** Anne Daebeler, Craig Herbold, Julia Vierheilig, Christopher J. Sedlacek, Petra Pjevac, Mads Albersten, Rasmus H. Kirkegaard, José R. de la Torre, Holger Daims, Michael Wagner

**Affiliations:** Division of Microbial Ecology, Department of Microbiology and Ecosystem Science, Research Network “Chemistry meets Microbiology”, University of Vienna, Austria; Center for Microbial Communities, Department of Chemistry and Bioscience, Aalborg University, Denmark; Department of Biology, San Francisco State University, San Francisco, CA, USA

**Keywords:** AOA, thaumarchaea, thermophile, nitrification, hot spring, nirK, polymerase, anaerobic metabolism

## Abstract

Ammonia-oxidizing archaea (AOA) within the phylum *Thaumarchaea* are the only known aerobic ammonia oxidizers in geothermal environments. Although molecular data indicate the presence of phylogenetically diverse AOA from the *Nitrosocaldus* clade, group 1.1b and group 1.1a *Thaumarchaea* in terrestrial high-temperature habitats, only one enrichment culture of an AOA thriving above 50 °C has been reported and functionally analyzed. In this study, we physiologically and genomically characterized a novel *Thaumarchaeon* from the deep-branching *Nitrosocaldaceae* family of which we have obtained a high (∼85 %) enrichment from biofilm of an Icelandic hot spring (73 °C). This AOA, which we provisionally refer to as “*Candidatus* Nitrosocaldus islandicus”, is an obligately thermophilic, aerobic chemolithoautotrophic ammonia oxidizer, which stoichiometrically converts ammonia to nitrite at temperatures between 50 °C and 70 °C. *Ca.* N. islandicus encodes the expected repertoire of enzymes proposed to be required for archaeal ammonia oxidation, but unexpectedly lacks a *nirK* gene and also possesses no identifiable other enzyme for nitric oxide (NO) generation. Nevertheless, ammonia oxidation by this AOA appears to be NO-dependent as *Ca.* N. islandicus is, like all other tested AOA, inhibited by the addition of an NO scavenger. Furthermore, comparative genomics revealed that *Ca.* N. islandicus has the potential for aromatic amino acid fermentation as its genome encodes an indolepyruvate oxidoreductase *(iorAB)* as well as a type 3b hydrogenase, which are not present in any other sequenced AOA. A further surprising genomic feature of this thermophilic ammonia oxidizer is the absence of DNA polymerase D genes - one of the predominant replicative DNA polymerases in all other ammonia-oxidizing *Thaumarchaea.* Collectively, our findings suggest that metabolic versatility and DNA replication might differ substantially between obligately thermophilic and other AOA.

## Introduction

*Thaumarchaea* (Brochier-Armanet *et al.*, 2008) are among the most abundant archaeal organisms on Earth, and thrive in most oxic environments (Francis *et al.*, 2007; Erguder *et al.*, 2009; Schleper and Nicol, 2010; Bouskill et al. 2012; Prosser and Nicol, 2012; Stahl and de la Torre, 2012; Stieglmeier *et al,* 2014a), but have also been detected in anoxic systems (Molina *et al.*, 2010; Bouskill *et al.*, 2012; Buckles *et al.*, 2013; Beam *et al.*, 2014; Lin *et al.*, 2015). This phylum comprises ammonia-oxidizing archaea (AOA) and other archaeal taxa in which ammonia oxidation has not been demonstrated. All cultured members of the *Thaumarchaea* are AOA and grow by using ammonia, urea or cyanate as substrate (Palatinszky *et al.* 2015; Bayer *et al.* 2016; Sauder *et al.* 2017; Qin *et al.* 2017a), although *in situ* experiments suggest that certain members of this phylum capable of ammonia oxidation also possess other lifestyles (Mußmann *et al.* 2011; Sauder *et al.* 2017). In aquatic and terrestrial environments *Thaumarchaea* often co-occur with ammonia-oxidizing bacteria (AOB), and frequently outnumber them by orders of magnitude (Francis *et al.*, 2005; Leininger *et al.*, 2006; Mincer *et al.*, 2007; Adair and Schwarz, 2008; Abell *et al.*, 2010; Mußmann *et al.*, 2011; Zeglin *et al.*, 2011; Daebeler *et al.*, 2012). *Thaumarchaea* also inhabit extreme environments like terrestrial hot springs and other high temperature habitats, where AOB are not detectable (Weidler *et al.*, 2007; Reigstad *et al.*, 2008; Wang *et al.*, 2009; Zhao *et al.*, 2011; Chen *et al.*, 2016). In addition to the presence of *Thaumarchaea* in hot environments, high *in situ* nitrification rates (Reigstad *et al.*, 2008; Dodsworth *et al.*, 2011; Chen *et al.*, 2016) and transcription of genes involved in archaeal ammonia oxidation in several hot springs over 74 °C (Zhang *et al.*, 2008; Jiang *et al.*, 2010) support an important role of thermophilic AOA in these systems.

Despite their apparent importance for nitrogen cycling in a wide range of thermal habitats, only one thermophilic [on the basis of the definition by Stetter (1998) that thermophiles grow optimally above 50 °C] AOA species from an enrichment culture has been reported to date (de la Torre *et al.*, 2008; Qin *et al.*, 2017b) and was named *Candidatus (Ca.)* Nitrosocaldus yellowstonensis. In addition, several enrichment cultures and one pure culture of moderately thermophilic AOA, which are able to grow at 50 °C, but grow optimally only at temperatures below 50 °C, have been described (Hatzenpichler *et al.*, 2008; Lebedeva *et al.*, 2013; Palatinszky *et al.*, 2015). Therefore, our current knowledge on specific adaptations or metabolic capabilities of thermophilic AOA growing preferably at temperatures above 50 °C is very limited (Spang *et al.*, 2012).

In 16S rRNA and ammonia monooxygenase subunit A *(amoA)* gene trees *Ca*. Nitrosocaldus yellowstonensis branches most deeply among *Thaumarchaea* that possess ammonia monooxygenase (AMO) genes. In consequence, the *Nitrosocaldales* clade has been considered as being close to the evolutionary origin of *Thaumarchaea* encoding the genetic repertoire for ammonia oxidation (Spang *et al.* 2017, de la Torre *et al.*, 2008). However, since the genome sequence of *Ca*. N. yellowstonensis is not yet published, phylogenomic analysis to confirm an ancestral position of the *Nitrosocaldales* relative to other *Thaumarchaea* have been pending.

Here we report on the enrichment, phylogenomic analyses, and selected (putative) metabolic features of a novel, obligately thermophilic, AOA from the *Nitrosocaldales* clade obtained from a biofilm collected from an Icelandic hot (73 °C) spring. This organism, provisionally referred to as *Ca*. Nitrosocaldus islandicus, occupies a fundamentally different niche compared to other genomically characterized AOA as its ammonia-oxidizing activity is restricted to temperatures ranging from 50 °C to 70 °C.

## Materials and Methods

### Enrichment, cultivation, and physiological experiments

The enrichment of *Ca.* N. islandicus was initiated by inoculation of 40 ml sterile mineral medium (Koch *et al.*, 2015) containing 0.5 mM filter-sterilized NH_4_Cl with approximately 0.1 g of hot spring biofilm, which had been submerged in running water at the sampling site in a geothermal area in Graendalur valley, (64° 1’ 7” N, 21° 11’ 20” W) Iceland. At the sampling site, the spring had a pH of 6.5 and a temperature of 73 °C. The culture was initially incubated without agitation in 100 ml glass bottles in the dark at 60 °C and checked weekly for ammonium and nitrite content of the medium by using Nessler’s reagent (K_2_H_g_I_4_ – KOH solution; Sigma-Aldrich) and nitrite/nitrate test stripes (Merkoquant; Merck). Ammonium (1 mM NH_4_Cl) was replenished when completely consumed. At the same time pH was monitored by using pH test stripes (Machery-Nagel) and kept at pH 7-8 by titration with NaHCO_3_. When the pH dropped below 6 the enrichment culture ceased to oxidize ammonia, but activity was restored by readjusting the pH to between 7 and 8. The ammonium and nitrite concentrations were quantified photometrically (Kandeler and Gerber, 1988; Miranda *et al.*, 2001) using an Infinite 200 Pro spectrophotometer (Tecan Group AG). The microbial community composition of the enrichment was regularly monitored by fluorescence *in situ* hybridization (FISH) with 16S rRNA-targeted probes labeled with dyes Cy3, Cy5, or Fluos as described elsewhere (Daims *et al.*, 2005). Probes targeting most bacteria (EUB338 probe mix; Amann et al., 1990; Daims et al., 1999), most archaea (Arch915, Stahl and Amann, 1991) and most *Thaumarchaea* (Thaum726, Beam, 2015) were applied. All positive results were verified using the nonsense probe nonEUB338 (Wallner *et al.* 1993) labeled with the same dyes. Treatments with the macrolide antibiotic spiramycin (15 mg l^-1^), which partly retains its antibacterial activity at 60 °C (Zorraquinio *et al.*, 2011), were performed as described in Zhang *et al.* (2015) together with serial dilutions ranging from 10^-5^ to 10^-8^ to obtain a highly enriched (∼ 85 %) AOA culture that was used for further characterization.

Growth rates of *Ca.* N. islandicus were determined across a range of incubation temperatures (50 °C to 70 °C). Triplicate cultures (25 ml) and negative controls (cultures not supplied with ammonium or inoculated with autoclaved biomass) were incubated for ten days in 100 ml Schott bottles without agitation in the dark at the respective temperature. Samples from these experiments were either stored at −20 °C for subsequent qPCR analyses (150 μl) or centrifuged (21,000 × g, 15min, 18 °C) to remove cells and the supernatant was stored at −20 °C for chemical analysis (600 μl. qPCR analysis with primers CrenamoA19F (Leininger *et al,* 2006) and CrenamoA616R (Tourna *et al.*, 2008) targeting the archaeal *amoA* gene was otherwise performed as described in Pjevac *et al.* (2017) before the genome sequence of *Ca.* N. islandicus was available. However, subsequent analysis demonstrated that the employed qPCR primers contain mismatches to the *amoA* sequence of this AOA in the middle of the forward and reverse primer. The specific growth rate was calculated from log-linear plots of *amoA* gene abundance in individual cultures. In this analysis, three out of seven time points were interpolated through linear regression.

To test whether the NO-scavenger 2-phenyl-4,4,5,5,-tetramethylimidazoline-3-oxide-1-oxyl (PTIO; TCI, Germany) inhibits ammonia oxidation by *Ca.* N. islandicus, 40 ml aliquots of mineral medium containing 1 mM ammonium were inoculated with 10 % (v/v) of an exponential-phase culture and incubated in duplicates in the presence of 0, 33, and 100 μM PTIO, respectively. PTIO was dissolved in sterile mineral medium before addition to the cultures. The cultures not exposed to PTIO were supplemented with the same volume of sterile medium. The cultures were sampled (2 ml) at the beginning of the experiment and after 15 days of incubation. Nitrite and ammonium concentrations were measured as described above.

### DNA extraction, genome sequencing, and annotation

DNA from three replicate enrichment cultures containing *Ca.* N. islandicus as the only detectable ammonia oxidizer was extracted as described by Angel and Conrad (2013) and sequenced by Illumina HiSeq next generation sequencing (250 bp paired end reads). Since we did not obtain a complete genome with this approach we re-extracted genomic DNA from the enrichment at a later stage according to Zhou *et al.* (1996) yielding high molecular weight DNA. Genomic DNA was then sheared in a Covaris g-TUBE (Covaris, USA) at 9000 RPM for 2x 1 min. in an Eppendorf mini spin plus centrifuge (Eppendorf, DE). The DNA was run on a E-Gel^™^ EX 1 % agarose gel (ThemoFisher, USA) and small DNA fragments were removed by excising a band with a length of ∼8 kb. The DNA was purified from the gel cut using the ultraClean 15 DNA Purification Kit (Qiagen, USA). The DNA was prepared for sequencing using the “1D Low Input gDNA with PCR SQK-LSK108” protocol (Oxford Nanopore Technologies, UK) and sequenced on a FLO-MIN106 flowcell using the MinION MK1b (Oxford Nanopore Technologies, UK) following the manufacturers protocol using MinKNOW (v. 1.7.14). The nanopore reads were basecalled using Albacore (V. 2.0.1) (Oxford Nanopore Technologies, UK). The complete genome was assembled using a hybrid approach combining the data from the Illumina and nanopore sequencing with the hybrid assembler Unicycler (v. 0.4.1, Wick et al., 2017). The genome bins of the two contaminating organisms were assembled from the Nanopore reads using Miniasm (Li *et al.*, 2016) and polished twice with the Nanopore reads using Racon (Vaser *et al.*, 2017). No other microbe encoding genes indicative for ammonia-oxidation was identified in either of the two the metagenomes.

The complete genome of *Ca.* N. islandicus was uploaded to the MicroScope platform (Vallenet *et al.*, 2013) for automatic annotation, which was amended manually where necessary. The full genome sequence of *Ca.* N. islandicus has been deposited in GenBank (accession CP024014) and associated annotations are publicly available in MicroScope (*Candidatus* Nitrosocaldus islandicus strain 3F).

Protein-coding genes from the novel *Thaumarchaeon* were compared to those from 30 *Thaumarchaea* with available genomic data (Table S1) downloaded from NCBI. The coding sequences (CDS) with accession numbers from each genome, as downloaded from NCBI, were combined with additional CDS predictions made by Prodigal (Hyatt *et al.*, 2010) to account for variability in CDS predictions from different primary data providers and platforms. Predicted CDS from the novel *Thaumarchaeon* were aligned to CDS from reference genomes using blastp (Word_size=2, substitution matrix BLOSUM45). Genes were considered homologous only if the blastp alignment exceeded 50 *%* of the length of both query and subject sequences. CDS of *Ca.* N. islandicus that lacked any homologs in other *Thaumarchaea* were considered “unique”. Unique CDS of unknown function were searched for secretion signals and for predicted membrane-spanning domains of the encoded proteins using the Phobius web server (Käll *et al.*, 2007) and putative structures were determined using the Phyre2 web server (Kelley *et al.*, 2015). Homology to “Thaumarchaea-core” proteins was assessed by cross-referencing the blastp homology search to the proteins defined for *Ca.* Nitrosotalea devanaterra by Herbold *et al.* (2017).

### Phylogenetic analysis and habitat preference

For 16S rRNA and *amoA* gene-based phylogenetic analysis, the full-length 16S rRNA and *amoA* gene sequences of *Ca.* N. islandicus retrieved from the genome assembly were imported into the ARB software package (Ludwig *et al.*, 2004) together with other full length 16S rRNA or *amoA* gene sequences from cultivated AOA strains and aligned with the integrated ARB aligner with manual curation. 171 sequences from the *Aigarchaea* were included in the alignment and used as outgroup in the 16S rRNA gene phylogenetic analyses. For the *amoA* gene phylogenetic analyses no outgroup was selected. The 16S rRNA and *amoA* gene consensus trees were reconstructed using Maximum-Likelihood (ML; using the GTRGAMMA evolution model), Neighbour Joining (NJ) and Maximum Parsimony (MP) methods. For all calculations, a sequence filter considering only positions conserved in ≥50 % of all *thaumarchaeal* and *aigarchaeal* sequences was used, resulting in 2444 and 488 alignment positions for the 16S rRNA and *amoA* genes, respectively.

A Bayesian-inference phylogenomic tree was obtained using the automatically generated alignment of 34 concatenated universal marker genes (Table S2), which were identified by CheckM in Parks *et al.* (2015). This alignment was used as input for PhyloBayes (Lartillot *et al.*, 2009) with ten independent chains of 4,000 generations using the CAT-GTR model; 2,000 generations of each chain were discarded as burn-in, the remainder were subsampled every second tree (bpcomp -x 2000 2 4000) and pooled together for calculation of posterior probabilities.

Whole-genome based average nucleotide identity (gANI, Varghese *et al.*, 2015) and average amino acid identity values (AAI, Konstantinidis and Tiedje, 2005) were calculated between the genomes of *Ca.* N. islandicus and *Ca.* N. yellowstonensis using sets of annotated genes supplemented with additional gene calls predicted by Prodigal (Hyatt *et al.*, 2010). gANI was calculated using the Microbial Species Identifier (MiSI) method (Varghese *et al.*, 2015). For AAI, bidirectional best hits were identified using blastp, requiring that query genes aligned over at least 70 *%* of their length to target genes (in each unidirectional blastp search). Query gene length was used to calculate a weighted average % identity over all best hit pairs and the calculations were repeated using each genome as query and target.

The occurrence of organisms closely related to *Ca.* N. islandicus and *Ca.* N. yellowstonensis in publicly deposited amplicon sequencing data sets was assessed using IMNGS (Lagkouvardos *et al.*, 2016) with the full-length 16S rRNA gene sequences of both organisms as query and a nucleotide identity threshold of 97 %.

*PolB* amino acid sequences were extracted from the arCOG database (arCOG14 ftp://ftp.ncbi.nih.gov/pub/wolf/COGs/arCOG/ (arCOG15272, arC0G00329, arC0G00328, arC0G04926, arC0G15270). Additional *thaumarchaeal polB* sequences were identified using *Ca.* N. islandicus as a query in a blastp search against the nr protein database. These additional *thaumarchaeal* sequences, the *polB* sequence from *Ca.* N. islandicus and arCOG database sequences were de-replicated using usearch (Edgar, 2010) with -sortbylength and -cluster_smallmem (-id 0.99 -query_cov 0.9), aligned using default settings in mafft (Katoh and Standley, 2013) and a phylogenetic tree was calculated using FastTree (Price *et al.*, 2010).

Nitrilase superfamily amino acid sequences were obtained from Pace and Brenner (2001). Alignment and phylogenetic reconstruction was carried out with Bali-Phy (Suchard and Redelings, 2006; randomize alignment, iterations=11000, burnin=6000). Posterior tree pools from 10 independent runs were combined to generate a 50 % PP consensus tree and to assess bipartition support.

A dataset for assessing the phylogenetic relationship of the alpha subunit of 2-oxoacid:ferredoxin oxidoreductases (OFORs) was based on Gibson *et al.* (2016) and supplemented with additional indolepyruvate oxidoreductase (ior) sequences. Genomes available (as of October 30, 2017) from the NCBI genomes database were downloaded, genes were predicted using Prodigal V2.6.3 (Hyatt *et al.*, 2010) and predicted genes were screened for *iorA* (TIGRFAM03336) using hmmsearch v3.1b2 (hmmer.org) with an e-value cutoff of 0.001. Genes meeting the search criteria were used as queries against the complete TIGRFAM database to ensure that the extracted *iorA* sequences matched the *iorA* model as the best-hit model with an e-value cutoff of 0.001. Reciprocal best-hit genes were required to align to the hmm over at least 500 contiguous bases. Amino acid sequences were then clustered into centroids using usearch v8.0.1517 (sortbylength and cluster_smallmem -id 0. 8 -query_cov 0.9; Edgar, 2010). Centroids were aligned using mafft v7.245 (Katoh and Standley, 2013) and trees were constructed using FastTree 2.1.4 (Price *et al.*, 2010). The initial phylogenetic placement of *Ca.* N. islandicus *iorA* in the resulting large tree (3,179 sequences) was used to choose a small set of bacterial *iorA* sequences to include in the final tree. The final dataset was aligned using mafft v7.245 (Katoh and Standley, 2013) and trees were constructed using FastTree 2.1.4 (Price *et al.*, 2010).

### Electron microscopy

For scanning electron microscopy, *Ca.* N. islandicus cells were harvested by centrifugation (4,500 × g, 15 min, 25 °C) and fixed on poly-L-lysine coated slides with a filter-sterilized 2.5 % glutaraldehyde fixation solution in phosphate buffered saline (PBS; 130 mM NaCl in 5 % [v/v] phosphate buffer mixture [20 to 80 v/v] of 200 mM NaH_2_PO_4_ and 200 mM Na_2_HPO_4_). Subsequently, fixed cells were washed three times for 10 min in PBS and post-fixed with a 1 % OsO4 solution in PBS for 40 min. The fixed cells were again washed three times in PBS, dehydrated in a 30 to 100 % (v/v) ethanol series, washed in acetone, and critical point dried with a CPD 300 unit (Leica). Samples were mounted on stubs, sputter coated with gold using a sputter coater JFC-2300HR (JEOL), and images were obtained with a JSM-IT300 scanning electron microscope (JEOL).

## Results and Discussion

### Enrichment and basic physiology of Ca. N. islandicus

An ammonia-oxidizing enrichment culture was established from biofilm material sampled from a hot spring located in the geothermal valley Graendalur of South-Western Iceland. Temperature tests for optimal activity and growth were performed at different time points during the enrichment period and showed varying results, but below 50 °C and above 75 °C activity and growth was never observed. Only during the initial enrichment phase did ammonia oxidation occur at 75 °C. At 65 °C the highest ammonia oxidation rates and the shortest lag phases were usually measured (data not shown), however in a single experiment the optimal temperature was 70 °C (Fig. S1). Likely, these variations reflect varying abundance ratios of *Ca.* N. islandicus and accompanying bacteria over time as described in Lebedeva *et al.* (2008). A high enrichment level of a single AOA phylotype (see below) was achieved by applying the antibiotic spiramycin (15 mg l^-1^) followed by biomass transfers into fresh medium using serial dilutions. This enrichment culture showed near stoichiometric conversion of ammonium to nitrite when incubated at 65 °C (Fig. 1). This was accompanied by growth of the AOA with a specific growth rate of 0.128 ± 0.011 d^-1^ (mean generation time of 2.32 ± 0.24 d), which is substantially slower than those reported for *Ca.* Nitrosocaldus yellowstonensis HL72, *N. viennensis* EN76, or *N. maritimus* SCM1 (de al Tore *et al.*, 2008; Könneke *et al.*, 2005; Martens-Habbena *et al.*, 2009; Stieglmeier *et al.*, 2014b; Table 1), but faster than a marine enrichment culture (Berg *et al.*, 2015).

**Figure 1.**
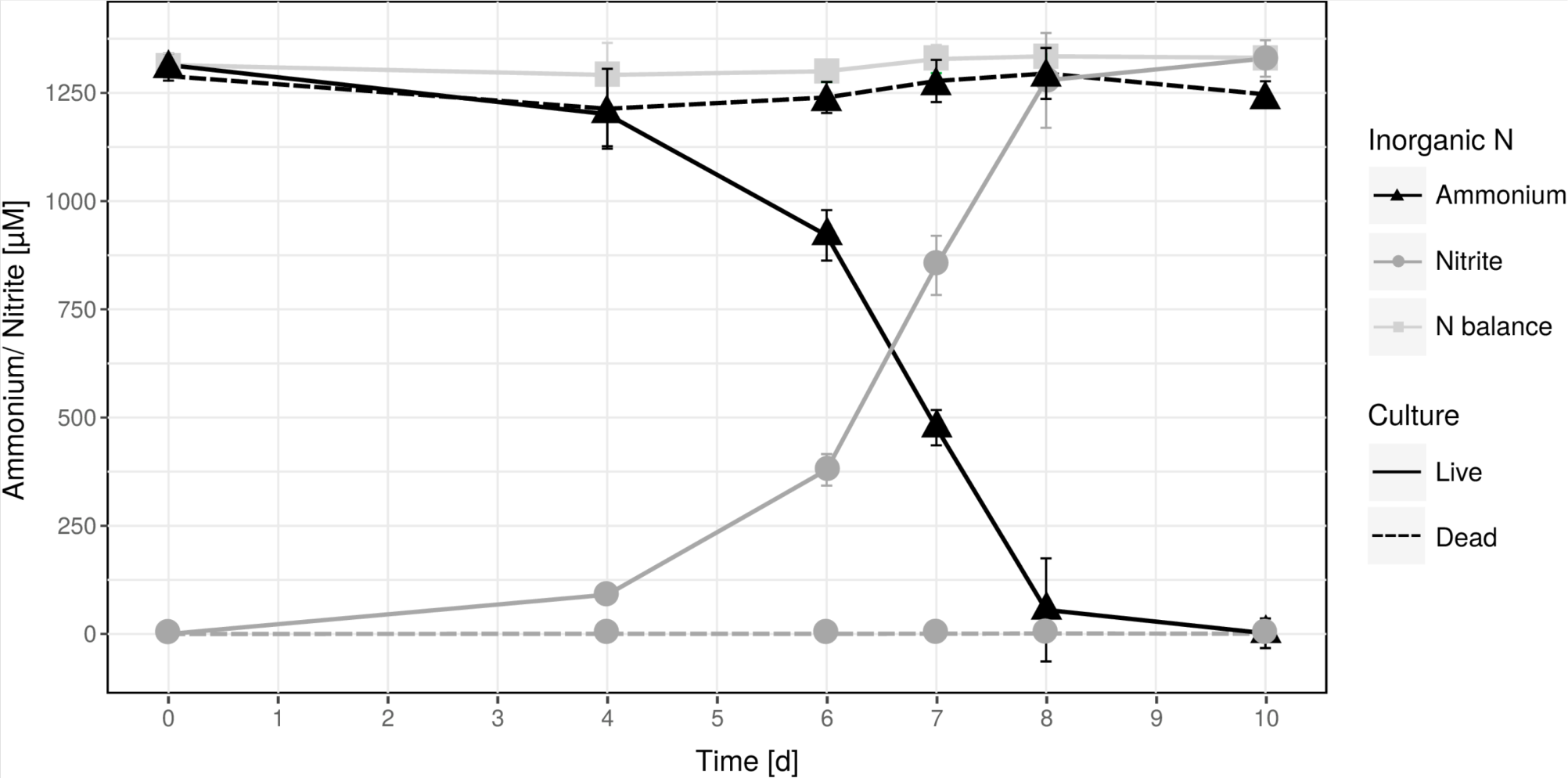
Near-stoichiometric oxidation of 1.25 mM ammonium to nitrite by the *Ca.* N. islandicus enrichment culture at 65 °C. Data points show means, error bars show 1 s.d. of n = 3 biological replicates. Solid and dashed lines denote live and dead culture incubations, respectively. If not visible, error bars are smaller than symbols.

**Table 1.**
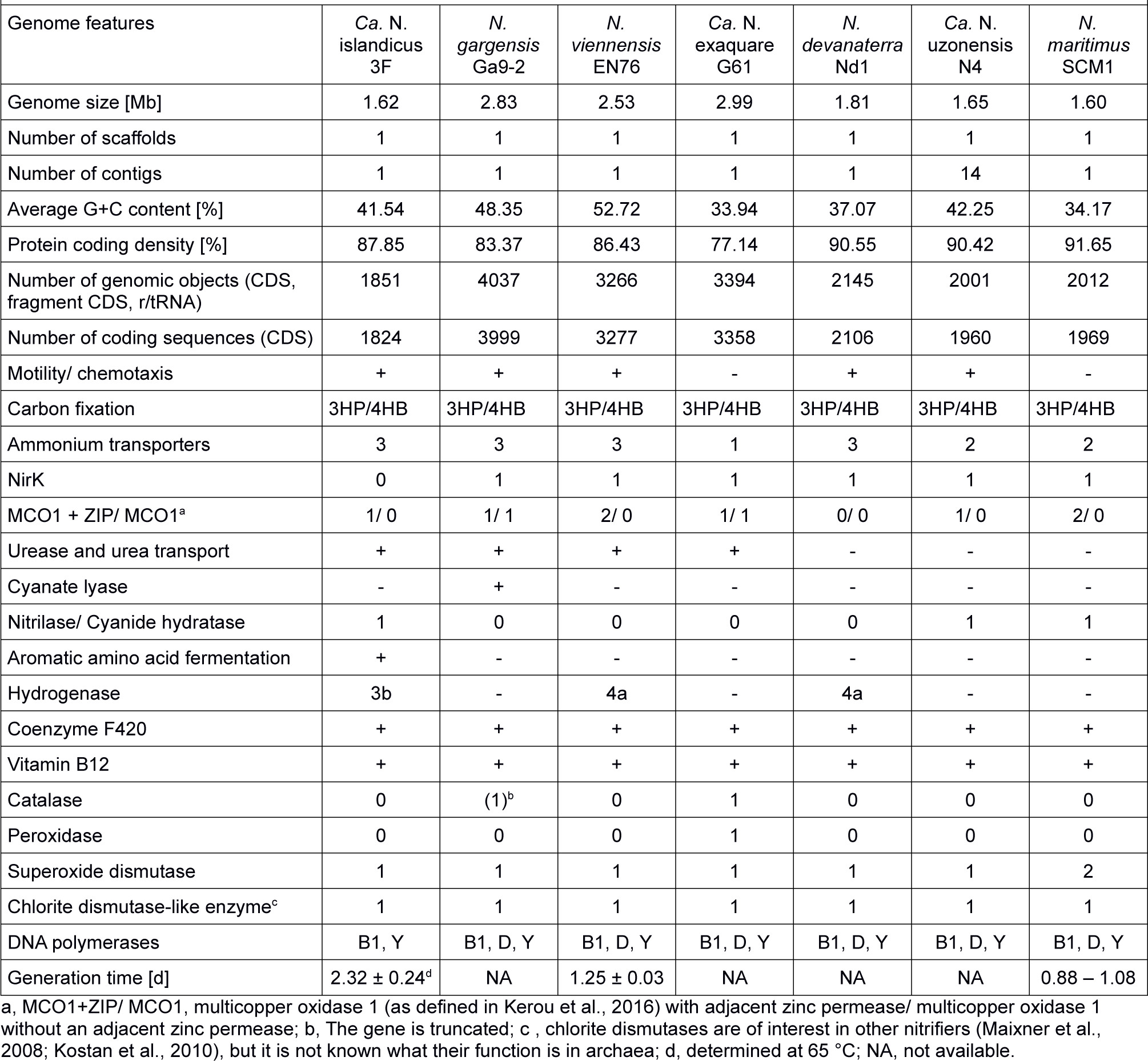
Genome features and growth rates of Ca. N. islandicus and of selected reference AOA

### Genome reconstruction, phylogeny, and environmental distribution

Metagenomic sequencing of the enrichment culture with Illumina and Nanopore demonstrated that the current culture contained an AOA as the only taxon encoding the repertoire genes required for ammonia oxidation. Hybrid assembly allowed reconstruction of the complete genome of this AOA as one circular contiguous sequence of 1.62 Mbps length (Table 1). The 16S rRNA gene and *amoA* gene of the newly enriched AOA are 96 and 85 % identical respectively to the genes of *Ca.* Nitrosocaldus yellowstonensis, the only other cultured obligately thermophilic AOA. The average amino acid sequence identity (AAI) and the genomic average nucleotide identity (gANI) between the genome and the one of *Ca.* N. yellowstonensis are 65.4 % (alignment fraction: 0.86) and 75.8 % (alignment fraction: 0.59), respectively, which is above the proposed genus and below the proposed species boundary thresholds (Qin *et al.*, 2014; Varghese *et al.*, 2015). Consequently, the enriched obligately thermophilic AOA was assigned to the same genus and referred to as *Ca.* Nitrosocaldus islandicus. According to 16S rRNA gene- and *amoA* gene-based phylogenies, *Ca.* N. islandicus is a novel member of the *Nitrosocaldales* clade, which seems to predominantly encompass AOA from thermal environments (Fig. 2). An extended phylogenomic analysis using a concatenated alignment of 34 proteins (Table S2) identified by CheckM (Parks *et al.*, 2015) confirmed that *Ca.* N. islandicus represents a basal lineage within the known ammonia-oxidizing *Thaumarchaea* (Fig. 3). This result lends strong support to the earlier notion, which was based on single-gene 16S rRNA and *amoA* phylogenies (de la Torre *et al.*, 2008), that the thermophilic *Nitrosocaldales* clade is an early diverging group of the ammonia-oxidizing *Thaumarchaea.* It would also be compatible with the possibility that archaeal ammonia oxidation originated in thermal environments (de la Torre *et al.*, 2008; Hatzenpichler *et al.*, 2008; Groussin *et al.*, 2011; Brochier-Armanet *et al.*, 2012).

**Figure 2.**
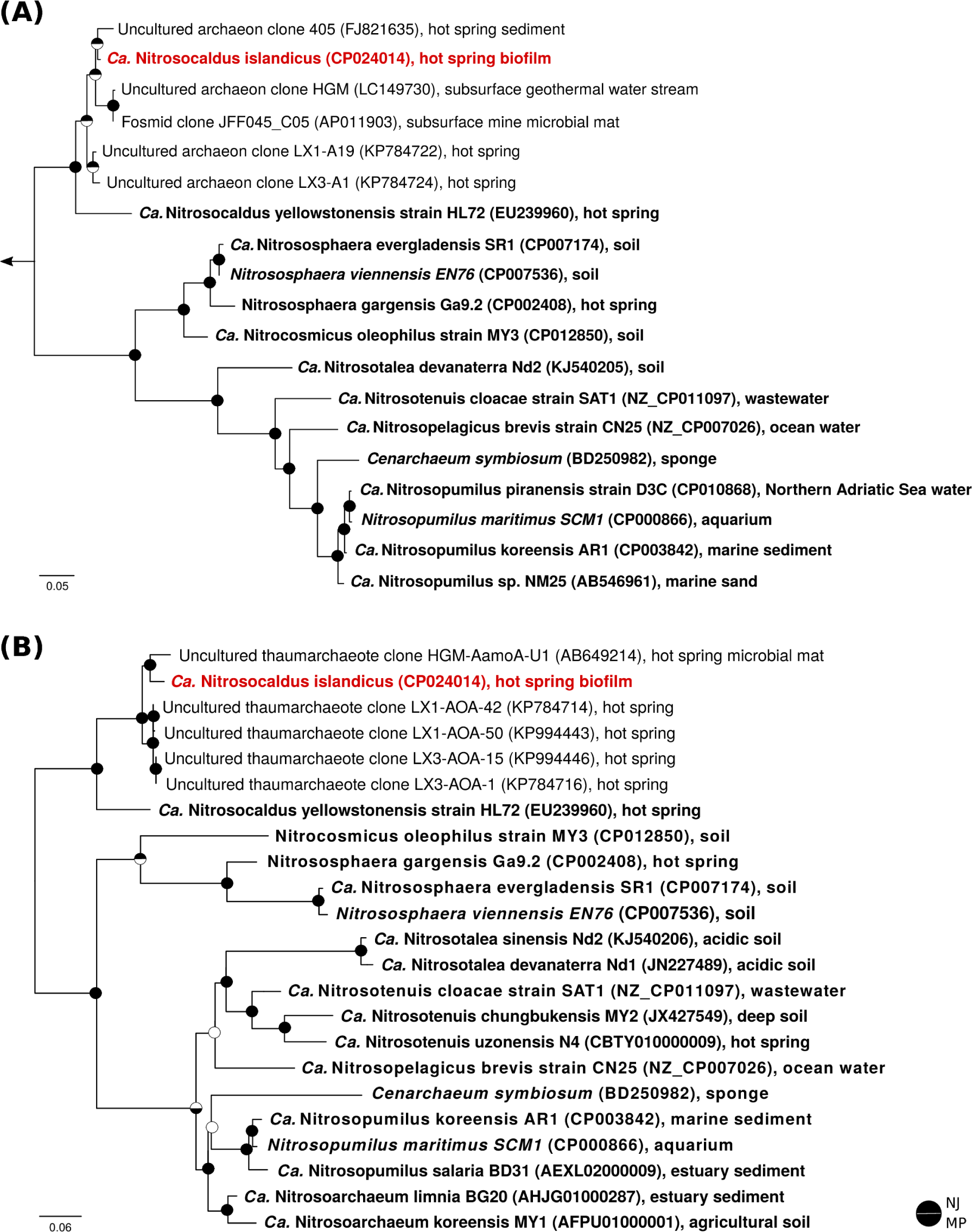
16S rRNA gene-based (A) and *amoA* gene-based (B) maximum likelihood phylogenies of representative *thaumarchaeal* sequences. For each sequence, the accession number and environmental source are indicated. Sequences from pure and enrichment cultures are depicted in bold, and *Ca.* N. islandicus is highlighted in red. The outgroup for the 16S rRNA tree were *aigarchaeal* sequences; the *amoA* phylogeny was calculated unrooted, but artificially rooted to the *Nitrosocaldales* afterwards. Circles at nodes denote support (filled) or no support (open) from Neighbour Joining (NJ, top half) and Maximum Parsimony (MP, bottom half) trees. The scale bars in panels (A) and (B) indicate 9 and 6 % sequence divergence, respectively.

**Figure 3.**
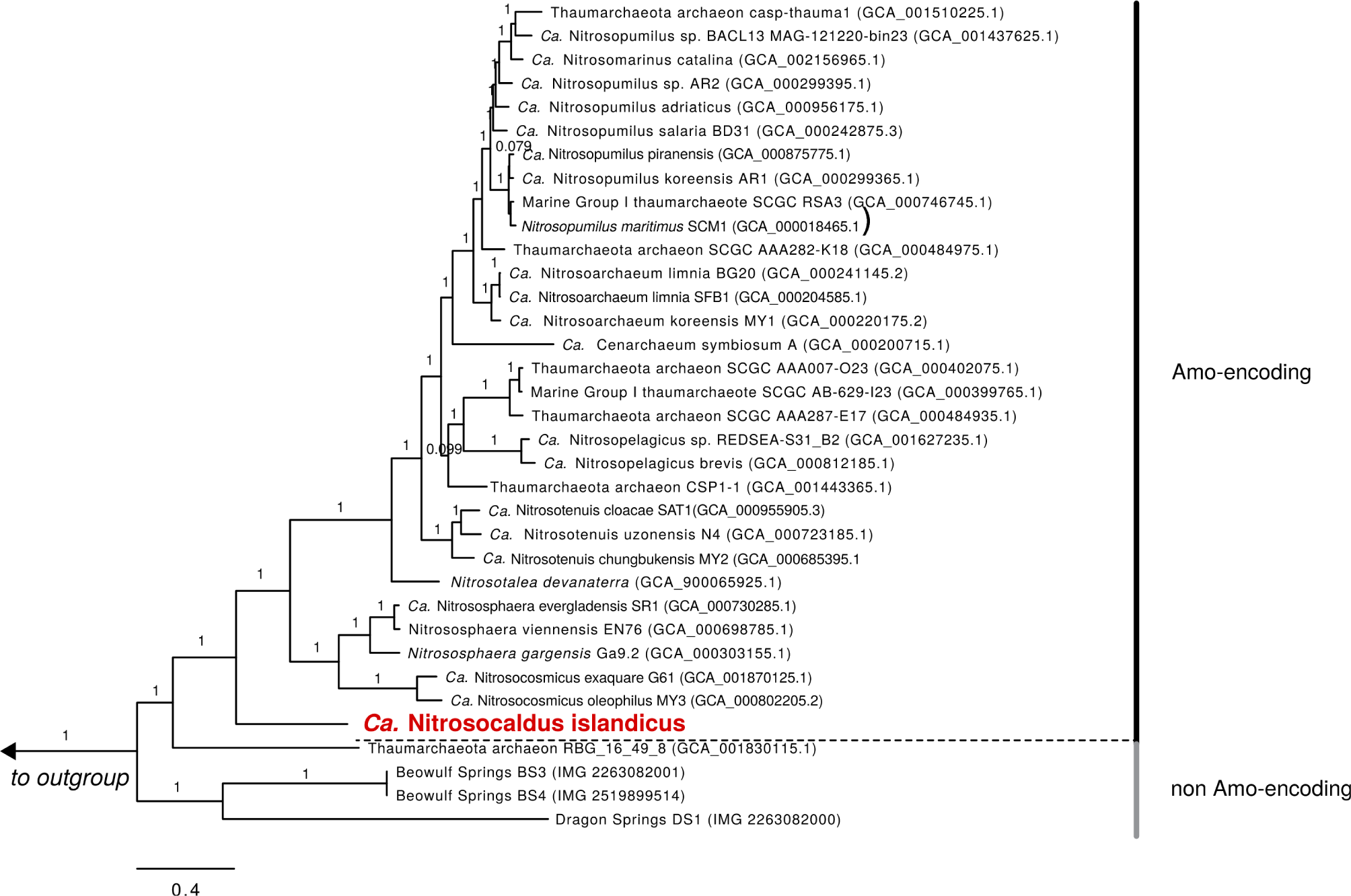
Bayesian inference tree of 34 concatenated universal marker proteins from 31 *amoA-* encoding *Thaumarchaea* including the *Nitrosocaldus-like* AOA and 4 non *amoA-*-encoding *Thaumarchaea-like* Archaea. Nineteen additional TACK-superphylum (Guy and Ettema, 2011) members (not shown) were used as an outgroup: *Aigarchaea* (assemblies GCA_000494145.1, GCA_000270325.1), *Bathyarchaea* (GCA_001399805.1, GCA_001399795.1, GCA_001593865.1, GCA_001593855.1, GCA_001593935.1, GCA_002011035.1, GCA_001273385.1), *Crenarchaea* (GCA_000011205.1, GCA_000591035.1, GCA_000253055.1, GCA_000813245.1), *Geothermarchaea* (GCA_002011075.1), *Korarchaea* (GCA_000019605.1), *Thorarchaea* (GCA_001563335.1, GCA_001563325.1), and *Verstraetearchaea* (GCA_001717035.1, GCA_001717015.1). Branches are labelled with average Bayesian posterior probability support over ten independent chains and the scale bar indicates 0.4 amino acid substitutions per site.

Metagenomic sequencing revealed that in addition to *Ca.* N. islandicus the culture also contained two heterotrophic bacterial contaminants, which were identified as a *Thermus sp.* and a member of the *Chloroflexi* phylum (Fig. 4). The enrichment level of *Ca.* N. islandicus was approximately 85 % based on read counts from the Nanopore sequencing, whereas the *Thermus sp.* and *Chloroflexi* accounted for 12 % and 3 %, respectively. FISH-analysis of the enrichment culture confirmed the dominance of *Ca.* N. islandicus and showed that the AOA grew mainly in aggregates, whereas the bacterial cells grew either co-localized with the archaeal flocs or planktonic (Fig. 5A). Electron microscopy demonstrated that the cells of *Ca.* N. islandicus are small (with a diameter of approximately 0.5 to 0.7 μm) and have an irregular coccoid shape (Fig. 5B). Morphologically they resemble the cells of *Ca.* N. yellowstonensis (Qin *et al.*, 2017b).

**Figure 4.**
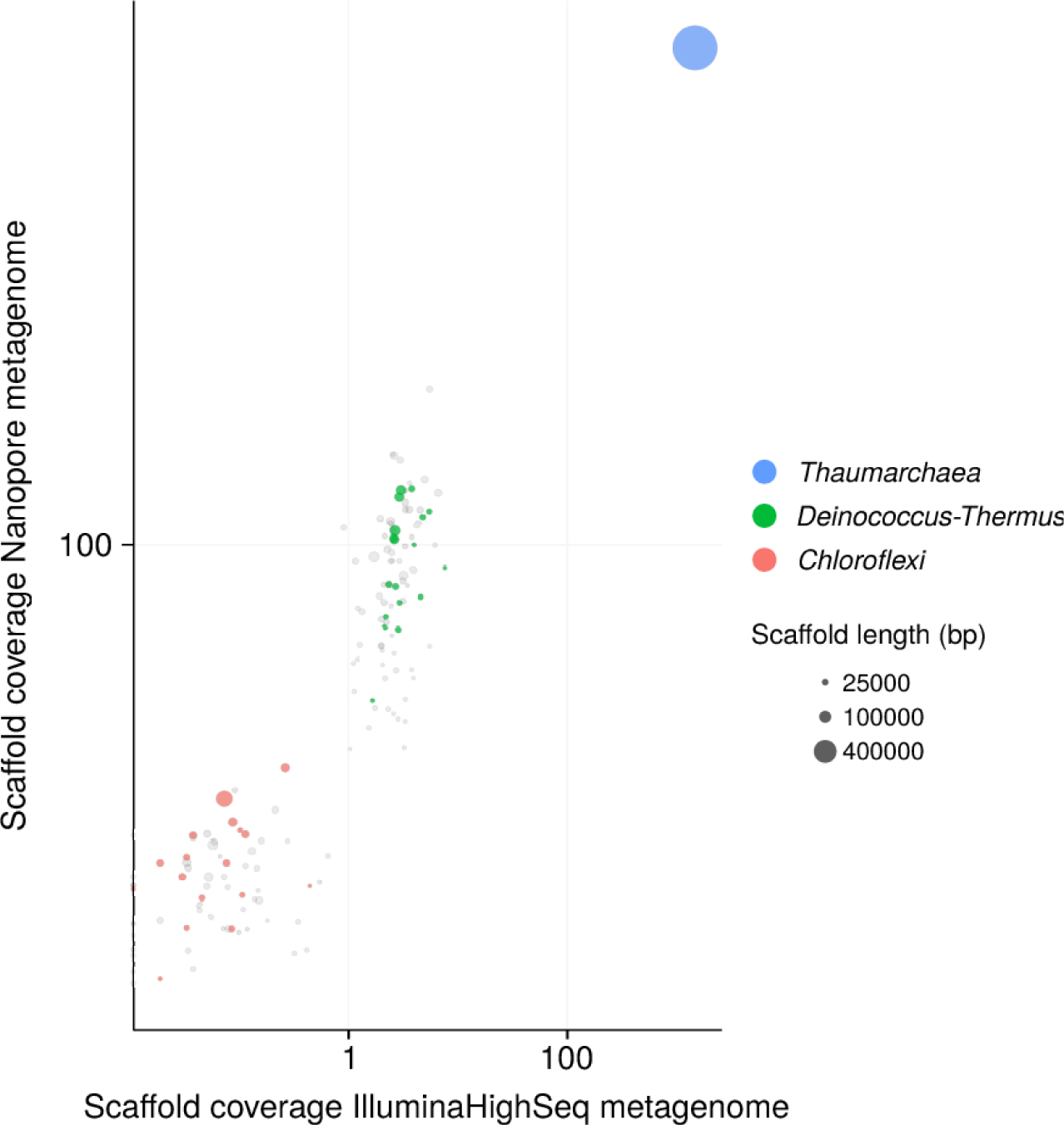
Sequence composition-independent binning of the metagenome scaffolds from two ammonia-oxidizing enrichment cultures. Circles represent scaffolds, scaled by the square root of their length. Clusters of similarly coloured circles represent potential genome bins. The x-axis shows binning of the scaffolds from an early enrichment culture, which still included other genera as well (not shown). The y-axis shows binning of the scaffolds from the latest enrichment culture containing only *Ca.* N. islandicus and the two remaining accompanying organisms. Genome bins for the *Thermus* (34 % complete) and the *Chloroflexi* (56 % complete) organism were obtained. The genome bin of the *Chloroflexi* organism contains genes that cluster within a clade of *Nitrobacter/ Nitrolancea* nitrite oxidoreductase *(nxrAB)* genes (data not shown). Since we did not observe nitrate production by the enrichment culture, the function of these genes remains unknown.

**Figure 5.**
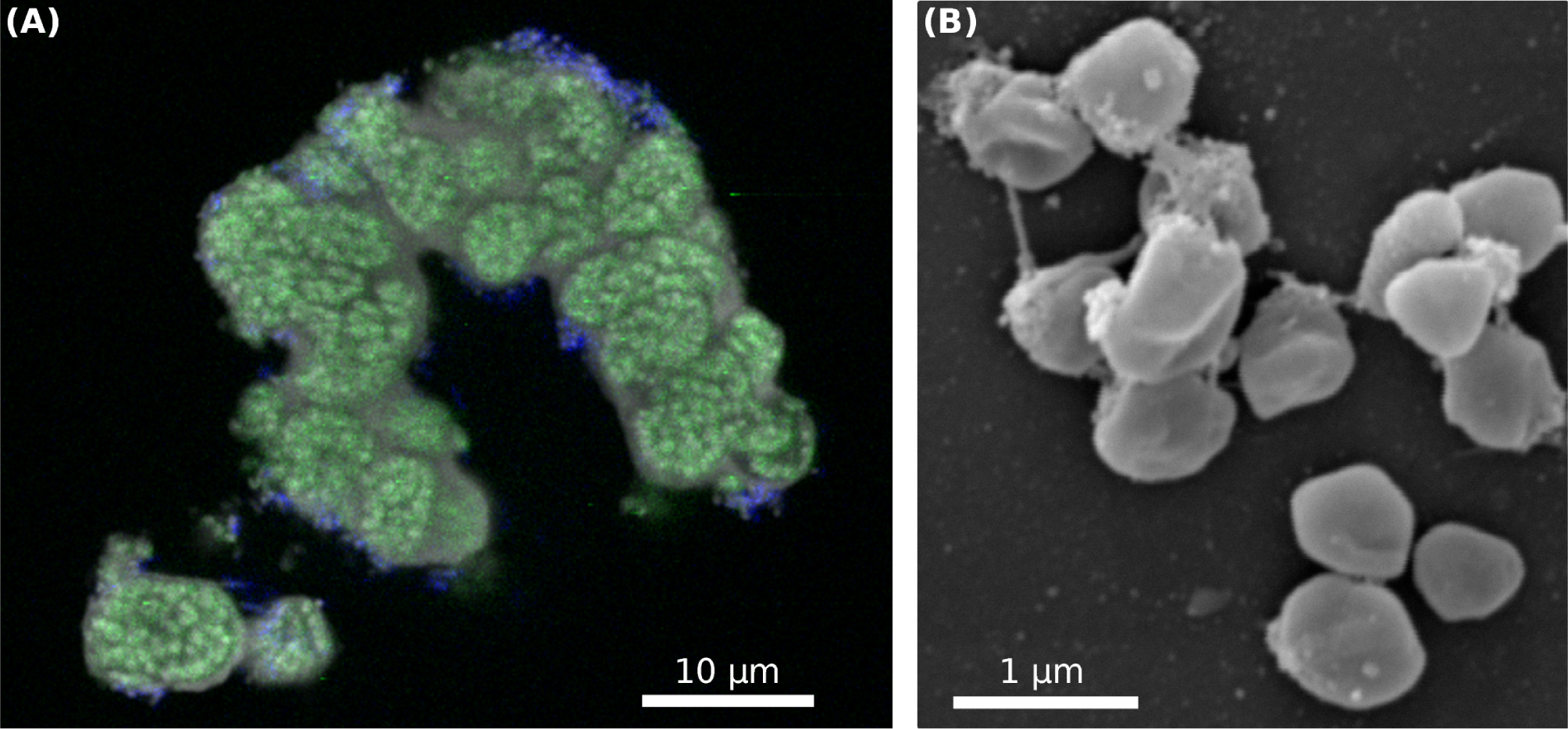
(A) FISH analysis of the enrichment culture illustrating the growth in microcolonies and the high relative abundance of *Ca.* N. islandicus. *Ca.* N. islandicus cells appear in green (stained by probe Thaum726 targeting most *Thaumarchaea)* and the bacterial contaminants in blue (labelled by probe EUB338). (B) Scanning electron micrograph of spherically shaped *Ca.* Nitrosocaldus islandicus cells. The cells have a diameter of 0.5 to 0.7 μm. *Ca.* N. islandicus cells were distinguishable from the rod-shaped bacterial contaminants by their smaller size and unique, ‘dented’ spherical shape.

The environmental distribution of the two cultured *Nitrosocaldales* members and closely related AOA was assessed by screening all publicly available 16S rRNA gene amplicon datasets (n=93,045) for sequences highly similar (97 %) to the 16S rRNA genes of *Ca.* N. islandicus and *Ca.* N. yellowstonensis using the pipeline described by Lagkouvardos *et al.* (2016). This analysis revealed that these taxa are highly confined in their distribution and occur predominantly in terrestrial hot springs where they can reach high relative abundances between 11.4 % and 86 % (*Ca.* N. islandicus and *Ca.* N. yellowstonensis, respectively) of the total microbial community (Fig. 6). Interestingly, *Ca.* N. yellowstonensis-related organisms seem to occur mainly in hot springs described as alkaline with a pH of around 8.5, but were also detected in a sample from a Tibetan wastewater treatment plant (Niu *et al.*, 2017). The unexpected detection of members of the *Nitrosocaldales* in the latter sample was confirmed by 16S rRNA-based phylogenetic analyses (data not shown) and it would be interesting to know whether this wastewater treatment plant is in some way connected to water from a close-by hot spring.

**Figure 6.**
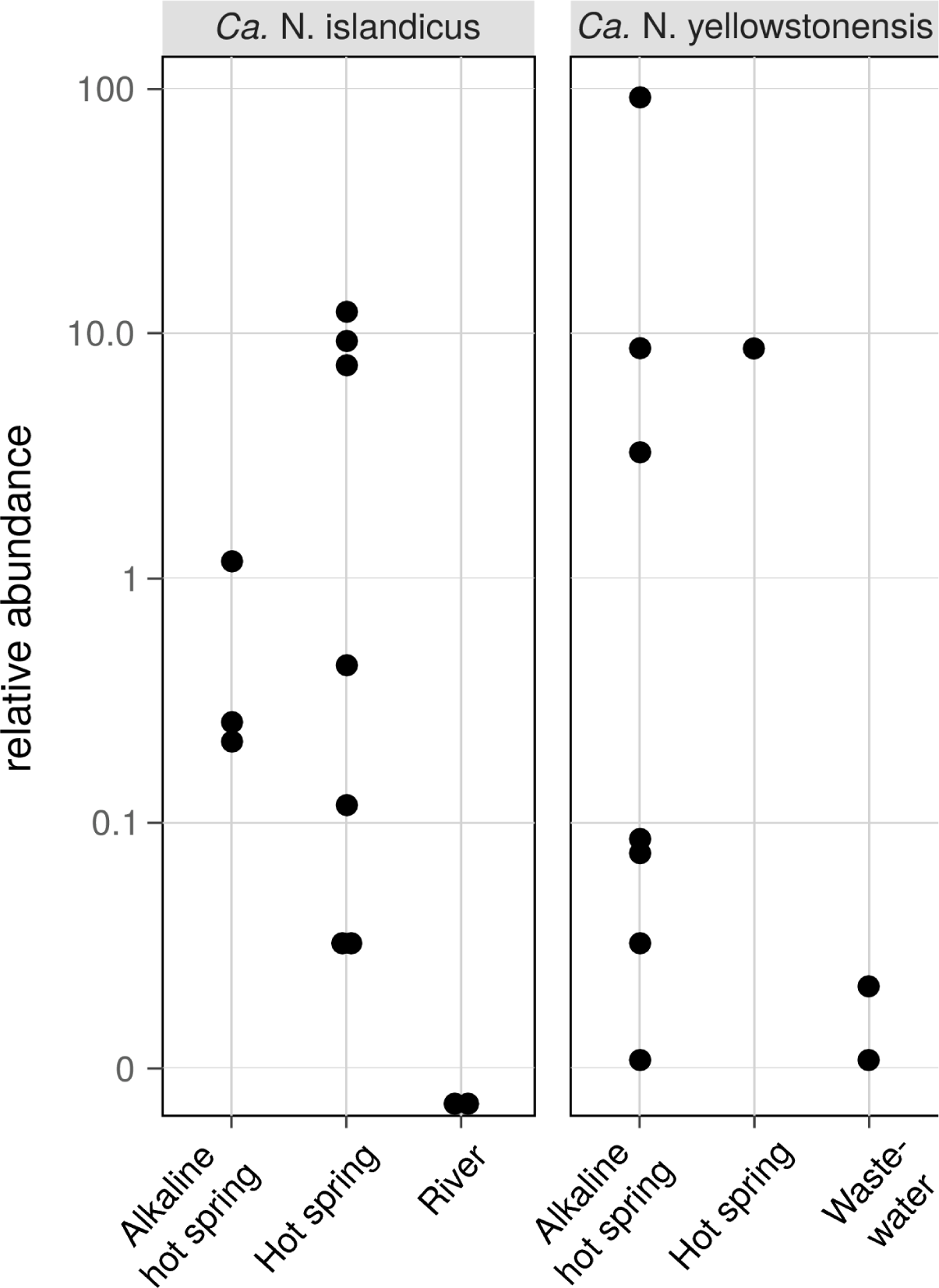
Occurrence and abundance of AOA related to *Ca.* Nitrosocaldus islandicus and *Ca.* Nitrosocaldus yellowstonensis in different habitats based on the presence of closely related 16S rRNA gene sequences in all public 16S rRNA gene amplicon datasets. Data shown are log-scale relative abundances of 16S rRNA gene sequences with a minimum similarity of 97 % in a sample (n=12 and n=10 out of 93,045 total datasets for *Ca.* N. islandicus and *Ca.* N. yellowstonensis, respectively). Sequences of the Tibetean waste water data set where retrieved from BioSample SAMN03464927 of Niu *et al.* (2017).

### Genome features

Addition of the complete genome of *Ca.* N. islandicus to the set of available thaumarchaeal genome sequences (n=30) reduced the number of gene families identified as representing the “Thaumarchaea-core” (Herbold *et al.*, 2017) from 743 to 669 (reduction by 9.96 %; Table S3). In a few cases, genes with low sequence homology to apparently absent core gene families are actually present in the genome of *Ca.* N. islandicus, but were not scored as they did not match the alignment length criterion. For example, *Ca.* N. islandicus, like all other AOA sequenced to date, has a gene encoding the K-subunit of RNA polymerase class I, but with a low sequence similarity to the respective orthologous genes in other AOA. In a few other cases, enzymes found in all other AOA genomes are absent but functionally replaced by members of different enzyme families. For example, all other genome-sequenced AOA contain a cobalamin-dependent methionine synthase. In contrast, *Ca.* N. islandicus possesses only an unrelated cobalamin-independent methionine synthase, which is also found in some other AOAs.

In addition to updating the *thaumarchaeal* core genome we also specifically looked for genes that are present in *Ca.* N. islandicus, but were not reported for other AOA before. In the following sections, the most interesting findings from these analyses are reported and put in context.

Like all other AOA, the *Ca.* N. islandicus genome encodes the typical repertoire for CO2 fixation via the modified 3-hydroxypropionate/4-hydroxybutyrate (3HP/4HB) cycle and for archaeal ammonia oxidation (Fig. 7; Fig. S2; Table 1) (Walker *et al.*, 2010; Spang *et al.*, 2012; Könneke *et al.*, 2014; Otte *et al.*, 2015; Kerou *et al.*, 2016). Unexpectedly however, the gene *nirK* encoding an NO-forming nitrite reductase is absent. NirK has been suggested to play an essential role for ammonia oxidation in AOA by providing NO for the NO-dependent dehydrogenation of hydroxylamine to nitrite (Kozlowski *et al.*, 2016). Interestingly, ammonia oxidation by *Ca.* N. islandicus was completely inhibited after the addition of ≥33 μM of the NO-scavenger PTIO (Fig. S3), a concentration that is lower or in the same range as previously reported to be inhibitory for other AOA (Shen et al., 2013; Jung et al., 2014; Martens-Habbena et al, 2015; Sauder et al., 2016).

**Figure 7.**
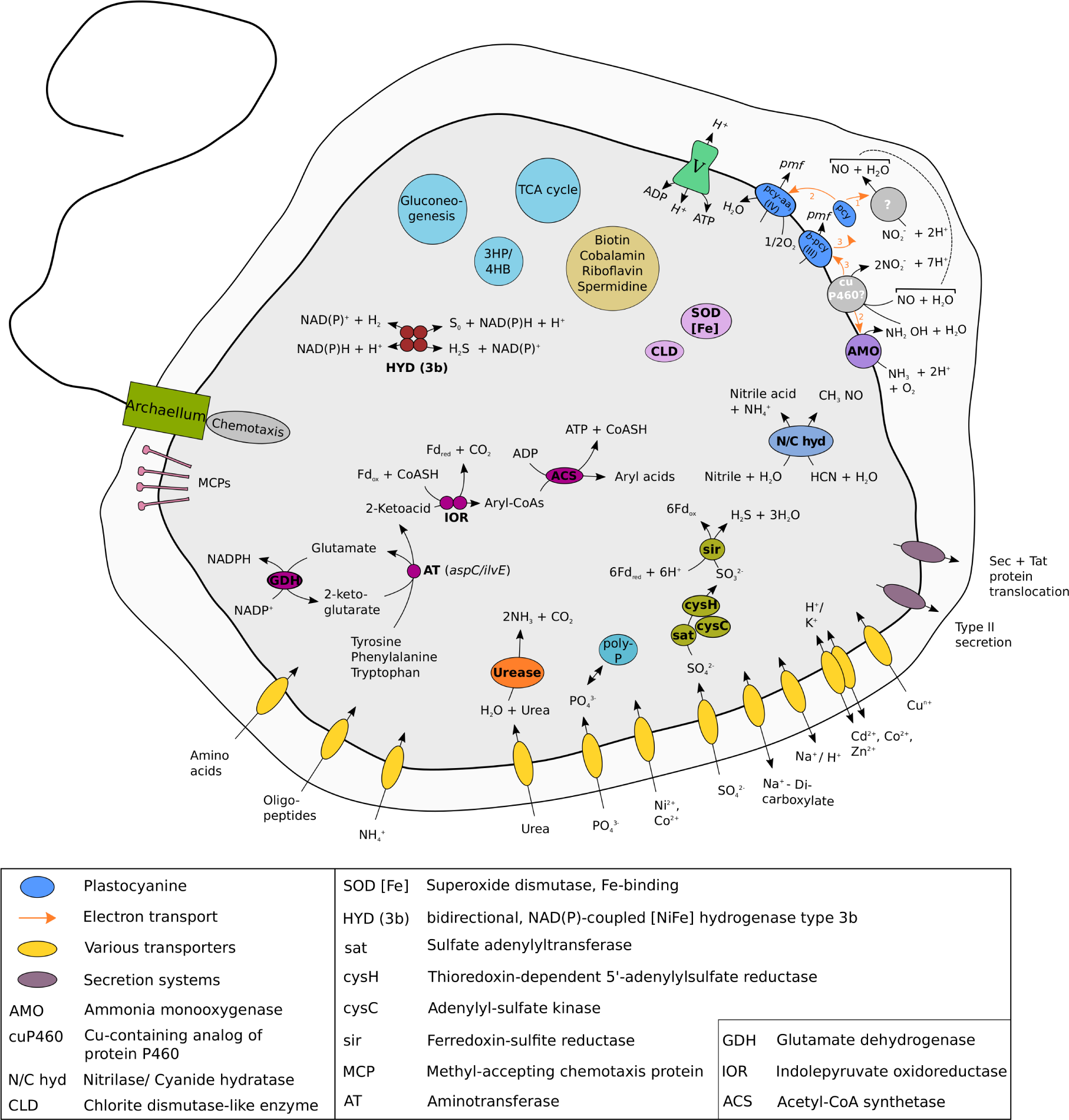
Cell metabolic cartoon constructed from the annotation of the *Ca.* N. islandicus genome. Enzyme complexes of the electron transport chain are labelled by Roman numerals. Most metabolic features displayed are discussed in the main text. The model of ammonia oxidation and electron transfer is depicted as proposed by Kozlowski *et al.* (2016). Locus tags of all genes discussed in the main text are given in table S4. Like some but not all AOA, *Ca.* N. islandicus encodes all genes required to assemble an archaeal flagellar apparatus that is composed of the flagellar filament, the motor, and its switch (Spang *et al.*, 2012; Lehtovirta-Morley *et al.*, 2016; Lebedeva *et al.*, 2013; Mosier *et al.*, 2012; Qin *et al.*, 2017; Li *et al.*, 2016), although no archaellum could be detected in our electron microscopic analysis. The *fla* gene cluster of *Ca.* N. islandicus shows a similar arrangement to *Nitrososphaera gargensis* and contains six genes including one gene for structural flagellin subunit FlaB/FlaA as well as the flagellar accessory genes *flaG*, *flaF*, *flaH, flaJ* and *flaI* (Spang *et al.*, 2012).

This finding suggests that NO is required for ammonia oxidation in *Ca.* N. islandicus despite the absence of NirK. The only other known AOA without a *nirK* gene are the sponge symbiont *Cenarchaeum symbiosum* (Hallam *et al.*, 2006; Bartossek *et al.*, 2010) and *Ca.* N. yellowstonensis (Stahl and de la Torre, 2012). For the uncultured *C. symbiosum* ammonia-oxidizing activity has not been demonstrated and the absence of *nirK* might have resulted from gene loss during adaptation to a life-style as symbiont. *Ca.* N. yellowstonensis is the closest cultured representative of *Ca.* N. islandicus, and the lack of *nirK* may thus be a common feature of the *Nitrosocaldales.* These AOA might produce NO by a yet unknown mechanism. In this context it is interesting to note that the hydroxylamine dehydrogenase of AOB, of which the functional homolog in archaea has not been identified yet, has recently been reported to produce NO instead of nitrite (Caranto and Lancaster, 2017). Alternatively, NO could be provided by accompanying organisms such as the *Thermus* and *Chloroflexi-like* bacteria that remain in the enrichment. Indeed, the genome bins obtained for these organisms both encode a *nirK* gene. The *Thermus sp.* genome bin further contains a *norBC* and *narGH* genes, in line with described denitrification capabilities for the genus *Thermus* (Alvarez *et al.*, 2014). A dependence of *Nitrosocaldales* on NO production by other microorganisms could explain why no pure culture from this lineage has been obtained yet.

*Ca.* N. islandicus possesses genes coding for urease that are present in some but not all AOA (Walker *et al.*, 2010; Spang *et al.*, 2012; Kerou *et al.*, 2016; Lehtovirta-Morley *et al.*, 2016; Sauder *et al.*, 2017) (Fig. 7; Table 1), but lacks a cyanase that is used by *Nitrososphaera gargensis* for cyanate-based growth (Palatinszky *et al.*, 2015). Additionally, the genome encodes an enzyme that either belongs to a novel class of the nitrilase superfamily or to the cyanide hydratase family (Fig. 7; Fig. S4). Nitrilases catalyze the direct cleavage of a nitrile to the corresponding acid while forming ammonia (Pace and Brenner, 2001) and cyanide hydratases convert HCN to formamide. Both substrates are relatively thermostable (Miyakawa *et al.*, 2002; Isidorov *et al.*, 1992). Nitriles occur as intermediates of microbial metabolism (Kobayashi *et al.*, 1993) and nitrile hydratases have previously been isolated from several thermophiles (Cramp *et al.*, 1997; Almatawah *et al.*, 1999; Kabaivanova *et al.*, 2008). Furthermore, both compounds are intermediates of the proposed abiotic synthesis of organics at hydrothermal sites (Miller and Urey, 1959; Schulte and Shock, 2005) and could thus be available in the hot spring habitat of *Ca.* N. islandicus. Similar genes have been found in the genomes of several other AOA from the *Nitrosopumilus* and *Nitrosotenuis* genera (Walker *et al.*, 2010; Mosier *et al.*, 2012; Lebedeva *et al.*, 2013; Park *et al.*, 2014; Bayer *et al.*, 2016) (Table 1) and it will be interesting to find out for which metabolism they may be used in AOA.

Intriguingly, *Ca.* N. islandicus might be able to ferment amino acids under anaerobic conditions as it contains the entire pathway used by some hyperthermophilic archaea for ATP generation from aromatic amino acids (Mai and Adams, 1994; Adams *et al.*, 2001; Ozawa *et al.*, 2012) (Fig. 7). In this pathway arylpyruvates are formed from aromatic amino acids by the activity of amino acid aminotransferases using 2-oxoglutarate as the amine group acceptor. The glutamate produced by this transamination can be recycled back to 2-oxoglutarate via glutamate dehydrogenase *(gdhA)* with the concomitant reduction of NADP+. With *ilvE* and *aspC* genes present, *Ca.* N. islandicus encodes at least two enzymes for which an aminotransferase activity specific for tyrosine, phenylalanine and aspartate has been demonstrated (Gelfand and Steinberg, 1977). Subsequently, these 2-ketoacids could be oxidatively decarboxylated and converted to aryl-CoAs by the oxygen-sensitive enzyme indolepyruvate oxidoreductase (Ozawa *et al.*, 2012) encoded by *iorAB* using oxidized ferredoxin as electron acceptor. *IorAB* is absent from all other genome-sequenced AOA and the *ior* genes present in *Ca.* N. islandicus have the highest similarity to and cluster together with iorAB-genes found in *Kyrpidia tusciae* and *Dadabacteria* (Fig. S5). Finally, transformation of aryl-CoAs to aryl acids catalyzed by ADP-dependent acetyl-CoA/acyl-CoA synthetase (Glasemacher *et al.*, 1997) leads to ATP formation via substrate-level phosphorylation (Fig. 7). *Ca.* N. islandicus encodes four acetyl-CoA/ acyl-CoA synthetases, two of which are most similar to non-syntenous homologs of acetyl-CoA/ acyl-CoA synthetases found in other AOA. However, the third gene is absent in all other AOA to date and its most similar homologs are encoded by species of the peptidolysing thermophilic archaea *Thermoproteus* and *Sulfolobus* and the fourth is most similar to an acetyl-/ acyl-CoA synthetase found in members of the thermophilic *Bathyarchaea* and *Hadesarchaea*.

The fermentation of aromatic amino acids also requires regeneration of oxidized ferredoxin (reduced by IorAB) and NADP+ (reduced by glutamate dehydrogenase). However, no canonical ferredoxin:NADP+ oxidoreductase, or other enzymes (Buckel and Thauer, 2013) described to regenerate oxidized ferredoxin, are encoded in the genome of *Ca.* N. islandicus. It seems unlikely that the amount of ferredoxin oxidized by an encoded ferredoxin-dependent assimilatory sulfite/nitrite reductase (Fig. 7) would be sufficient to compensate for all ferredoxin reduced in the dissimilatory fermentation pathway. However, Ca. N. islandicus can also oxidize reduced ferredoxin with a 2:oxoglutarate-ferredoxin oxidoreductase (Fig. S5). NAD(P)H can be re-oxidized by a cytosolic, bidirectional, NAD(P)-coupled type 3b [NiFe] -hydrogenase that is encoded by *Ca.* N. islandicus in contrast to all other genomically characterized AOA (Fig. 7; Table 1). NAD(P)H oxidation by this hydrogenase could lead to hydrogen generation, or the enzyme could act as a sulfhydrogenase that reduces zero valent sulfur compounds (produced by other organisms or present in the environment) to hydrogen sulfide (Ma *et al.*, 1993, Adams *et al.*, 2001). The hydrogenase genes are clustered at a single locus and code for the four subunits of the holoenzyme and accessory proteins (Fig. S6). This hydrogenase might also allow *Ca.* N. islandicus to use hydrogen as energy source providing reduced NAD(P)H under oxic conditions as this type of hydrogenase has been shown to tolerate exposure to oxygen (Bryant and Adams, 1989; Berney *et al.*, 2014; Kwan *et al.*, 2015).

Surprisingly, the genome of *Ca.* N. islandicus lacks genes for both subunits of the DNA polymerase D (PolD), which is present in all other AOA and most archaeal lineages (including thermophiles) with the exception of the Crenarchaea (Cann *et al.*, 1998; Makarova *et al.*, 2014, Saw *et al.*, 2015) (Table 1). It is assumed that either PolD alone or together with DNA polymerases of the B family (PolB) is required for DNA synthesis and elongation in these archaea (Cubonová *et al.*, 2013; Ishino and Ishino, 2013; Makarova *et al.*, 2014). The *Ca.* N. islandicus genome encodes only one B-type DNA polymerase (PolB1, Fig. S7) and one DNA polymerase of the Y family (PolY), generally considered to be involved in the rescue of stalled replication forks and enhancement of cell survival upon DNA damage (Friedberg *et al.*, 2002). Recently, it has been demonstrated for the PolD-lacking Crenarchaeon *Sulfolobus acidocaldarius* that both its PolB1 and PolY have polymerase activities *in vitro* (Peng *et al.*, 2016). However, *Ca.* N. islandicus (like other AOA) does not encode the PolB1-binding proteins PBP1 and PBP2, which are required to form a multisubunit DNA polymerase holoenzyme together with PolB in the Crenarchaeon *Sulfolobus solfataricus* P2 (Yan *et al.*, 2017). We hypothesize that *Ca.* N. islandicus may utilize one or both of the present polymerases for DNA replication, possibly in combination with its heterodimer PriSL, which has been demonstrated to function as a primase, a terminal transferase and a polymerase capable of polymerizing RNA or DNA chains of up to 7,000 nucleotides (Lao-Sirieix and Bell, 2004).

It is also interesting to note that the obligate thermophile *Ca.* N. islandicus like all genome-sequenced *Thaumarchaea* (Spang *et al.*, 2017) does not encode a reverse gyrase, which is widespread in hyperthermophilic microbes including other archaea of the TACK superphylum (Heine and Chandra, 2009; Makarova *et al.*, 2007; López-García *et al.*, 2015), but is not essential for growth under these conditions (Atomi *et al.*, 2004).

In conclusion, we have obtained a highly enriched (∼ 85 %) culture of an obligately thermophilic AOA from a hot spring in Iceland. Despite the impressive diversity of AOA in high temperature environments as revealed by molecular tools (Zhang et al., 2008; Wang et al., 2009; Zhao et al., 2011; Nishizawa *et al.*, 2013; Li *et al.*, 2015; Chen *et al.*, 2016), cultivation of only a single obligately thermophilic AOA species - *Ca.* Nitrosocaldus yellowstonensis - was reported before (de la Torre *et al.*, 2008). The newly enriched AOA represents a new species of the genus *Nitrosocaldus* and was named *Ca.* N. islandicus. Comparative analysis of its closed genome revealed several surprising features like the absence of DNA polymerase D and the lack of canonical NO-generating enzymes, although physiological experiments with a NO-scavenger demonstrated NO-dependent ammonia-oxidation, as described for other AOA (Shen *et al.*, 2013; Jung *et al.*, 2014; Martens-Habbena *et al.*, 2015; Sauder *et al.*, 2016). Furthermore, *Ca.* N. islandicus encodes the enzymatic repertoire for fermentation of aromatic amino acids that is, so far, unique among sequenced AOA. A pure culture of *Ca.* N. islandicus will be required to physiologically verify this genome-based hypothesis. Peptide or aromatic amino acid fermentation would enable an anaerobic lifestyle of *Ca.* N. islandicus and, if more widespread among *Thaumarchaea* not yet characterized (including mesophiles), might help explain their sometimes surprisingly high abundance in anaerobic ecosystems (Molina *et al.*, 2010; Bouskill *et al.*, 2012; Buckles *et al.*, 2013; Beam *et al.*, 2014; Lin *et al.*, 2015).

Based on the data presented here, we propose the following provisional taxonomic assignment for the novel *Thaumarchaeon* in our enrichment culture.

Nitrosocaldales order

Nitrosocaldaceae fam.

‘*Candidatus* Nitrosocaldus islandicus’ sp. nov.

#### Etymology

Nitrosus (Latin masculine adjective): nitrous; caldus (Latin masculine adjective): hot; islandicus (Latin masculine genitive name): from Iceland. The name alludes to the physiology of the organism (ammonia oxidizer, thermophilic) and the habitat from which it was recovered.

#### Locality

The biofilm of a terrestrial hot spring in Graendalur geothermal valley, Iceland (64° 1’7” N, 21° 11’20” W)

#### Diagnosis

An obligately thermophilic, aerobic chemolithoautotrophic ammonia oxidizer from the phylum *Thaumarchaea* growing as small irregular shaped cocci.

## Acknowledgements

We thank Bjarni Sigurdson and the Agricultural University of Iceland for permission to access and sample in Graendalur valley, Iceland. Daniela Gruber from the Core Facility of Cell Imaging and Ultrastructure Research at the University of Vienna and Stefano Romano are gratefully acknowledged for help with sample preparation for electron microscopy. The authors are grateful to Anja Spang for fruitful discussions. JV, CS, CH, and MW were supported by the ERC Advanced Grant NITRICARE (294343) to MW. AD and HD were supported by the Austrian Science Fund (FWF) grants P25231-B21 and T938.

## Author contributions

AD, JV and CS cultivated and enriched the culture; AD, CS and PP performed growth and activity experiments; AD performed FISH and SEM analysis; CH, AD, PP, MA and RK performed bioinformatic analysis; JT kindly provided access to the *Ca.* N. yellowstonensis genome; AD, JV, CS, PP, MW and HD manually curated the annotation of the genome and interpreted the genome data; AD and MW wrote the manuscript with help from all co-authors.

## Supplemental information

**Table S1.**
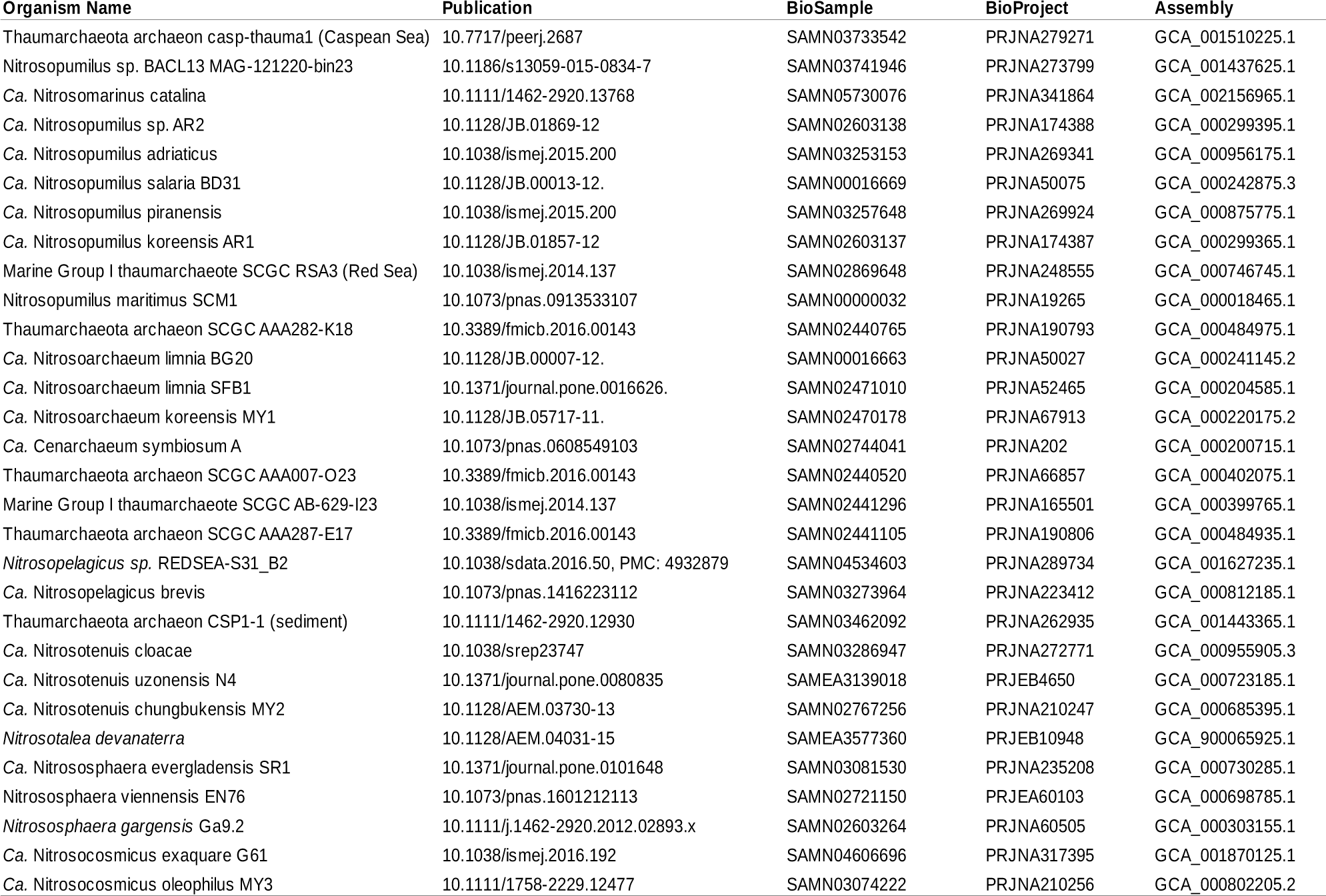
*Thaumarchaea* used for comparison of protein coding genes

**Table S2.** Marker genes used for phylogenomic tree

| Gene | Pfam Id | Length | Description |
| --- | --- | --- | --- |
| Alanine – tRNA ligase | TIGR00344 | 847 | Alanine – tRNA ligase |
| Ribosomal protein L10 | PF00466 | 100 | Ribosomal protein L10 |
| Ribosomal protein L11 | PF03946 | 60 | Ribosomal protein L11, N-terminal domain |
| Ribosomal protein L11 | PF00298 | 69 | Ribosomal protein L11, RNA binding domain |
| Ribosomal protein L13 | PF00572 | 128 | Ribosomal protein L13 |
| Ribosomal protein L14p/L23e | PF00238 | 122 | Ribosomal protein L14p/L23e |
| Ribosomal protein L16p/L10e | PF00252 | 133 | Ribosomal protein L16p/L10e |
| Ribosomal protein L18p/L5e | PF00861 | 119 | Ribosomal protein L18p/L5e |
| Ribosomal protein L1p/L10e | PF00687 | 220 | Ribosomal protein L1p/L10e |
| Ribosomal protein L22p/L17e | PF00237 | 105 | Ribosomal protein L22p/L17e |
| Ribosomal protein L23 | PF00276 | 92 | Ribosomal protein L23 |
| Ribosomal protein L29 | PF00831 | 58 | Ribosomal protein L29 |
| Ribosomal protein L3 | PF00297 | 263 | Ribosomal protein L3 |
| Ribosomal protein L4/L1 | PF00573 | 192 | Ribosomal protein L4/L1 |
| Ribosomal protein L5 | PF00281 | 56 | Ribosomal protein L5 |
| Ribosomal protein L5 | PF00673 | 95 | Ribosomal protein L5p, C-terminus |
| Ribosomal protein S11 | PF00411 | 110 | Ribosomal protein S11 |
| Ribosomal protein S12/S23 | PF00164 | 122 | Ribosomal protein S12/S23 |
| Ribosomal protein S15 | PF00312 | 83 | Ribosomal protein S15 |
| Ribosomal protein S17 | PF00366 | 69 | Ribosomal protein S17 |
| Ribosomal protein S19 | PF00203 | 81 | Ribosomal protein S19 |
| Ribosomal protein S2 | PF00318 | 211 | Ribosomal protein S2 |
| Ribosomal protein S3 | PF00189 | 85 | Ribosomal protein S3, C-terminal domain |
| Ribosomal protein S5 | PF03719 | 74 | Ribosomal protein S5, C-terminal domain |
| Ribosomal protein S5 | PF00333 | 67 | Ribosomal protein S5, N-terminal domain |
| Ribosomal protein S7p/S5e | PF00177 | 148 | Ribosomal protein S7p/S5e |
| Ribosomal protein S8 | PF00410 | 129 | Ribosomal protein S8 |
| Ribosomal protein S9/S16 | PF00380 | 121 | Ribosomal protein S9/S16 |
| Ribosomal Protein L2 | PF03947 | 130 | Ribosomal Proteins L2, C-terminal domain |
| Ribosomal protein L2 | PF00181 | 77 | Ribosomal proteins L2, RNA binding domain |
| RNA polymerase beta subunit | PF04563 | 203 | RNA polymerase beta subunit |
| RNA polymerase Rpb1 | PF04997 | 337 | RNA polymerase Rpb1, domain 1 |
| RNA polymerase Rpb1 | PF00623 | 166 | RNA polymerase Rpb1, domain 2 |
| RNA polymerase Rpb1 | PF05000 | 108 | RNA polymerase Rpb1, domain 4 |
| RNA polymerase Rpb2 | PF04561 | 190 | RNA polymerase Rpb2, domain 2 |
| RNA polymerase Rpb2 | PF04565 | 68 | RNA polymerase Rpb2, domain 3 |
| RNA polymerase Rpb2 | PF00562 | 386 | RNA polymerase Rpb2, domain 6 |
| RNA polymerase Rpb2 | PF04560 | 82 | RNA polymerase Rpb2, domain 7 |
| RNA polymerase Rpb6 | PF01192 | 57 | RNA polymerase Rpb6 |
| Signal peptide binding domain | PF02978 | 104 | Signal peptide binding domain |
| Translation-initiation factor 2 | PF11987 | 109 | Translation-initiation factor 2 |
| TruB family pseudouridylate synthase | PF01509 | 149 | TruB family pseudouridylate synthase |
| Valine – tRNA ligase | TIGR00422 | 863 | Valine – tRNA ligase |

**Table S3.**
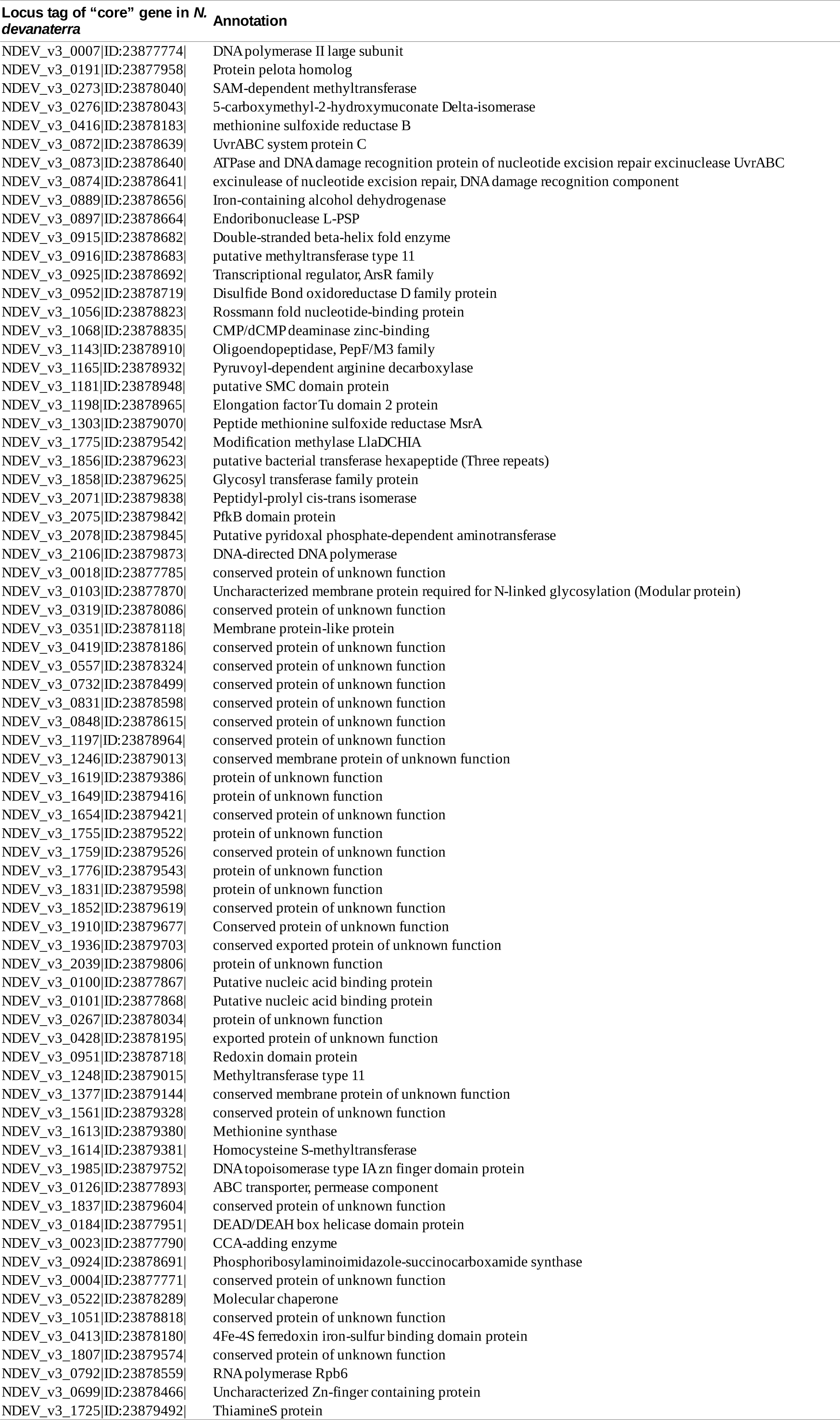
Genes not present in *Ca.* N. islandicus, but previously present in the “*Thaumarchaea-core*” as defined by^27^

**Table S4.**
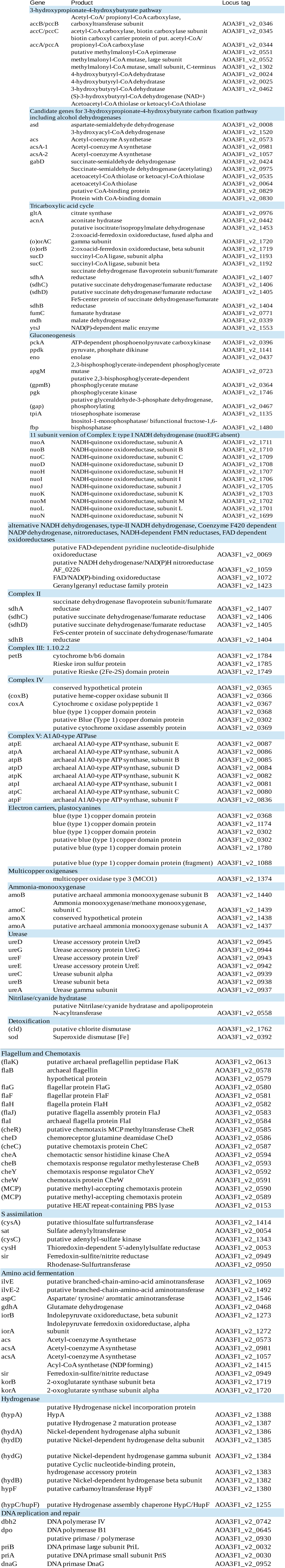
Genome locus tags and annotations of genes discussed in the main text.

**Figure S1.**
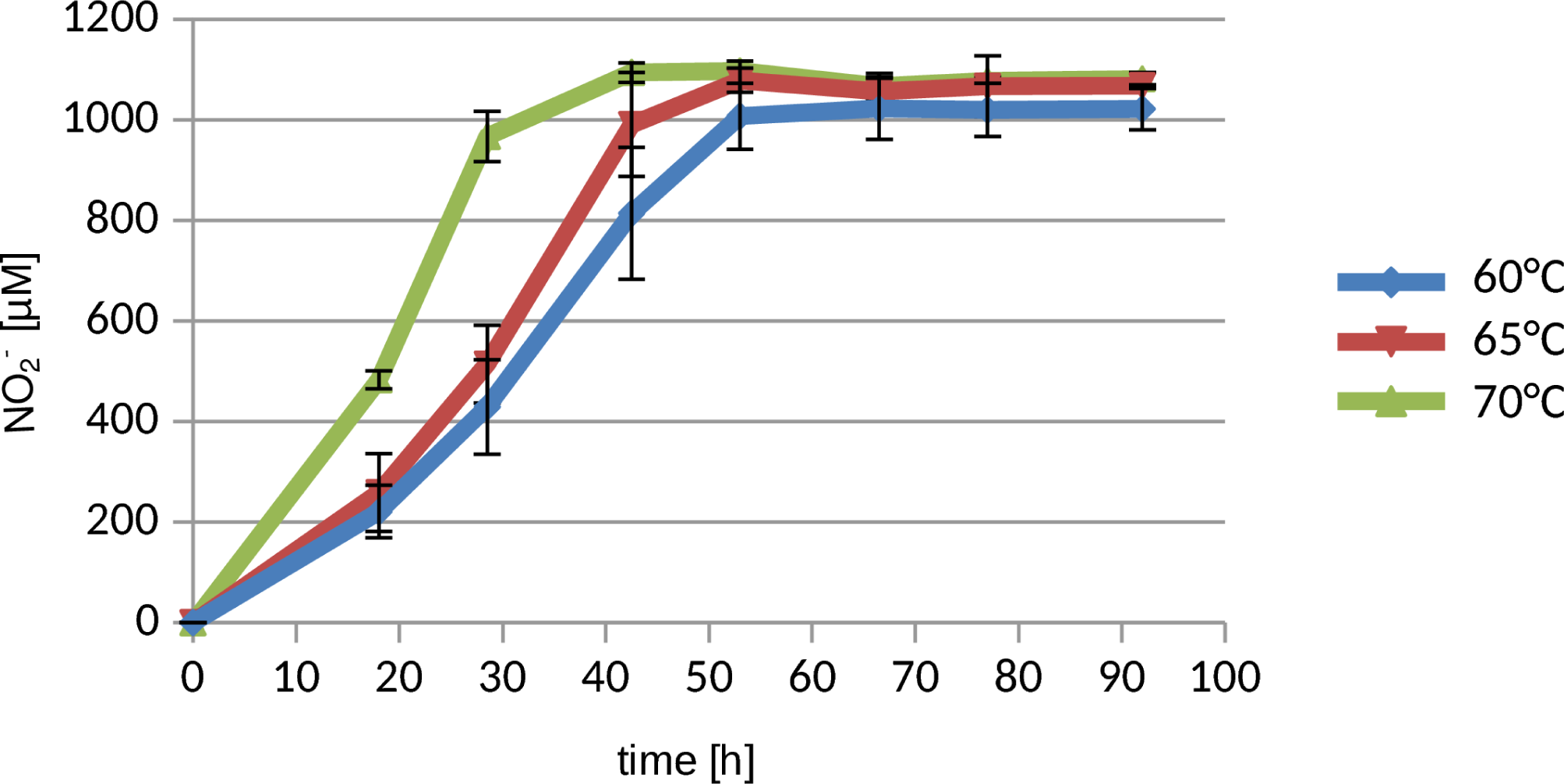
Nitrite accumulation through oxidation of ammonia at three different temperatures. Data points show means, error bars show standard errors of n = 3 biological replicates.

**Figure S2.**
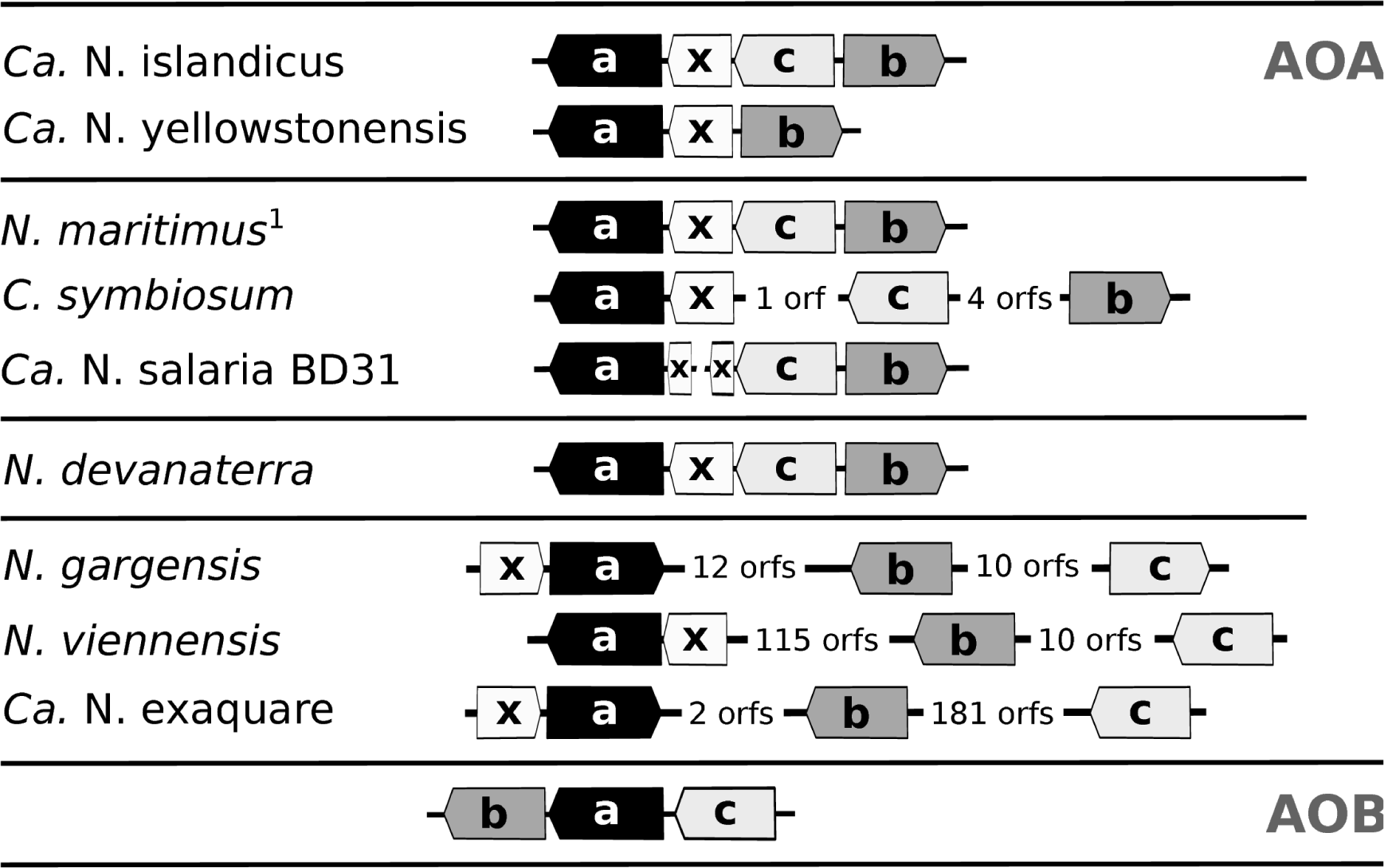
Gene order and orientation of the ammonia monooxygenase subunits (*amoA, amoB, amoC,* and the putative *“amoX”*) in *Ca.* N. islandicus and other Thaumarchaea. The gene order of ammonia-oxidizing bacteria (AOB) is given on the bottom as a reference. ^1^ also represents the gene arrangement in *Ca.* N. limnia, *Ca.* N. koreensis and *Ca.* N. uzonensis. The figure is a modified version of the figure 26.3 in *(28)*

**Figure S3.**
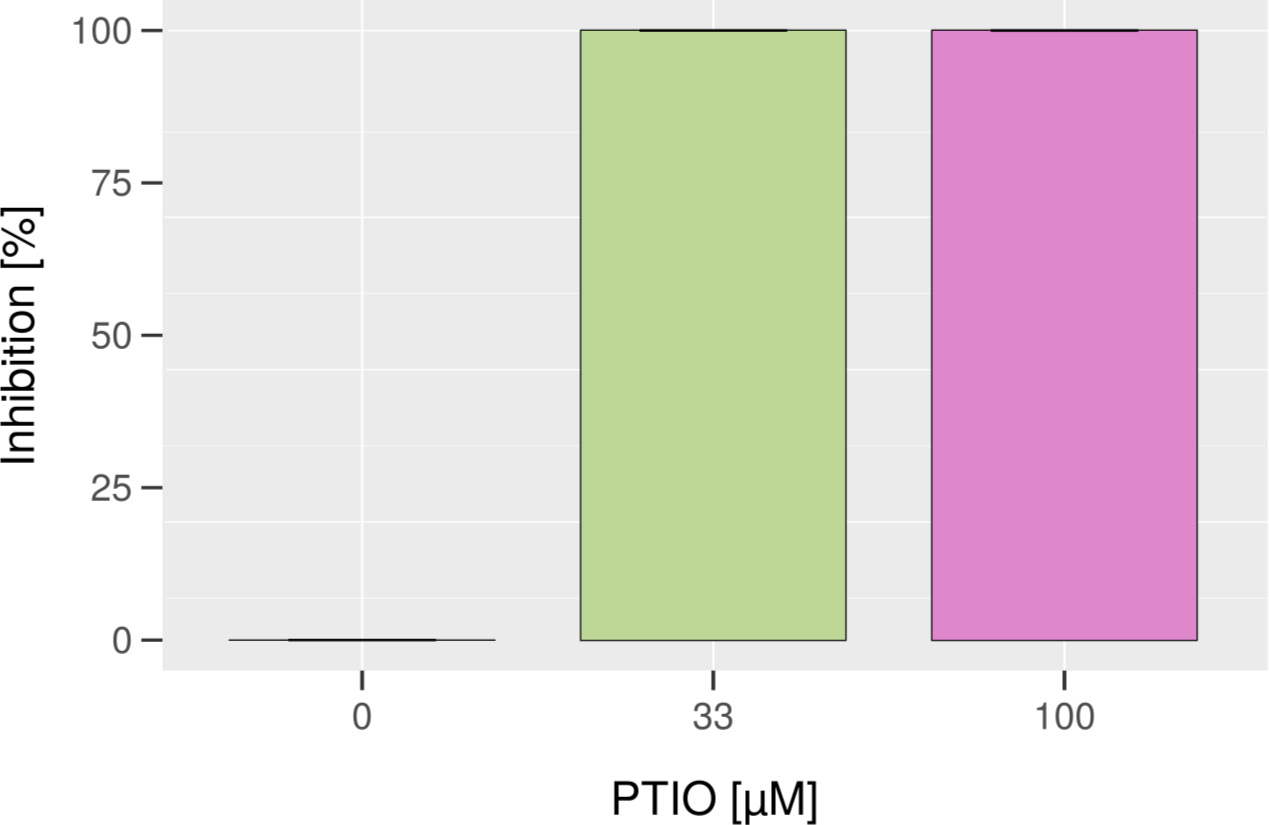
Inhibition of ammonia oxidation by *Ca.* N. islandicus caused by different concentrations of the NO-scavenger 2-phenyl-4,4,5,5-tetramethylimidazoline-1-oxyl 3-oxide (PTIO). Error bars show the standard error of two replicates.

**Figure S4.**
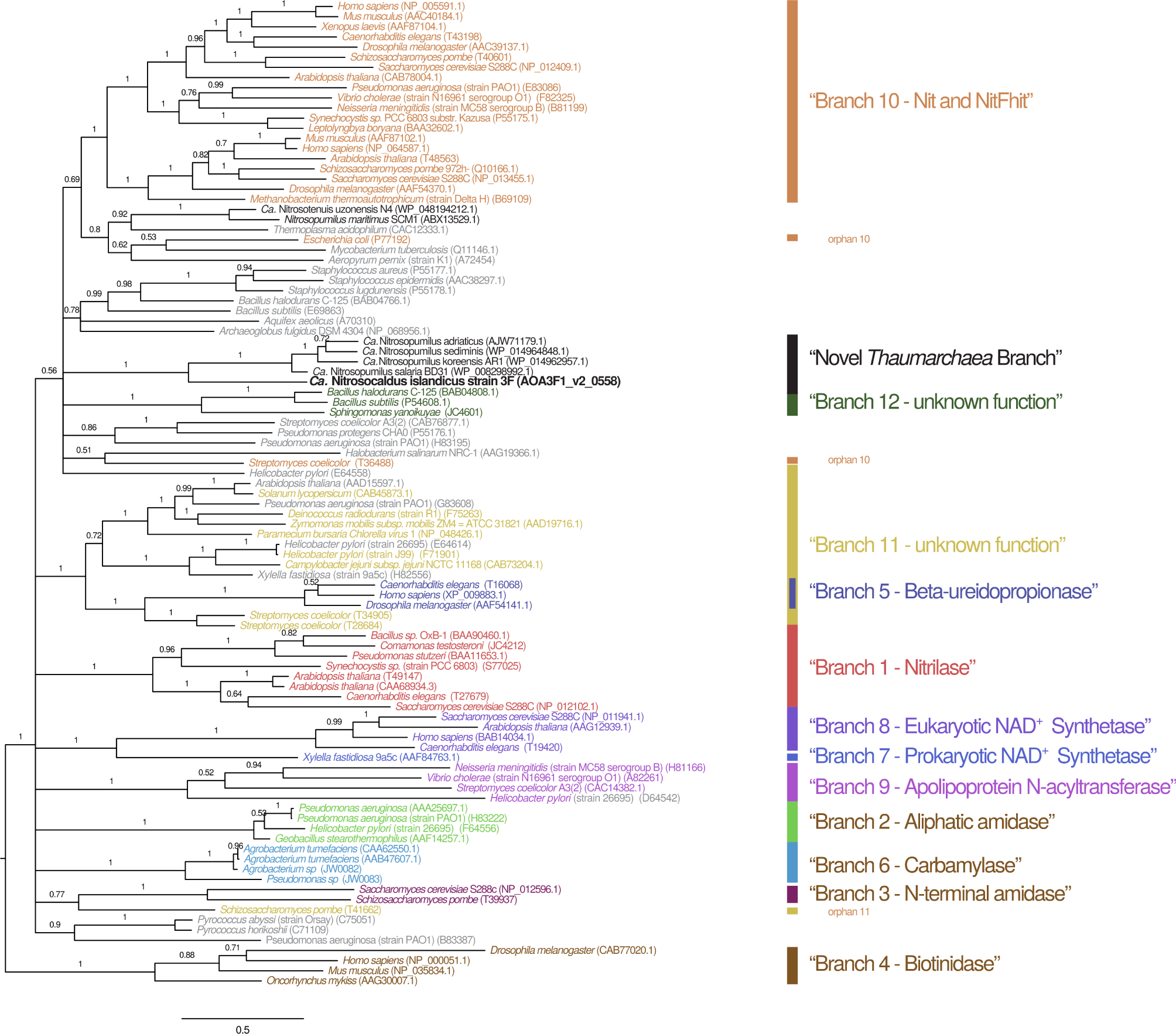
Phylogenetic tree of nitrilase superfamily members showing placement of *Ca.* Nitrosocaldus islandicus strain 3F in a novel branch consisting of *Thaumarchaea* (Novel *Thaumarchaea* branch). Sequences and branch labels are from (*29).* Grey labels indicate “nonfused outliers” as indicated by (29) which were not assigned to any of the 12 named branches. Black labels indicate sequences obtained from the *thaumarchaeal* genomes. Sequences assigned to branches 10 and 11 in (29) that do not clade with other members of those branches in this phylogenetic tree are labelled “orphan”.

**Fig. S5.**
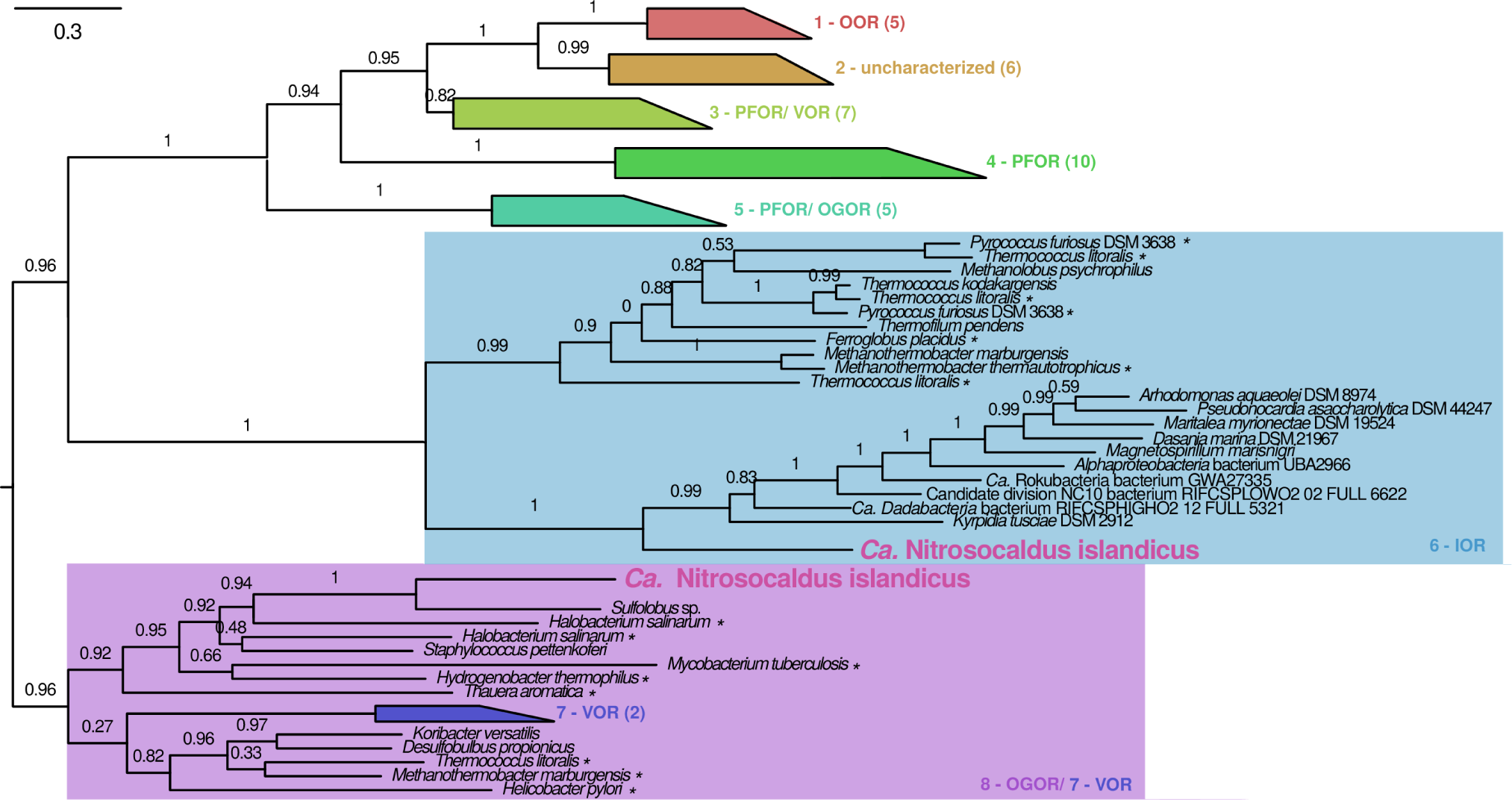
Unrooted phylogenetic tree of 2-oxoacid oxidoreductases (OFORs) showing placement of the two OFORs present in *Ca.* Nitrosocaldus islandicus strain 3F. Clade labels and most sequences are from *(30).* Numbers in brackets show the number of sequences in collapsed clades. Functionally characterized enzymes are indicated with a “*”. OOR, oxalate oxidoreductase; PFOR, pyruvate:ferredoxin oxidoreductase; VOR, 2-ketoisovalerate oxidoreductase; OGOR, 2-oxoglutarate oxidoreductase; IOR, indolepyruvate oxidoreductase.

**Figure S6.**
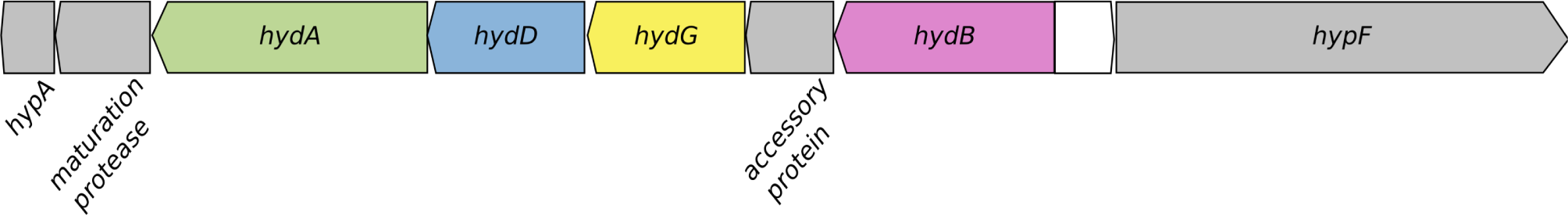
Schematic illustration of the genomic locus in *Ca.* Nitrosocaldus islandicus that encodes a bidirectional, NADP(H)-coupled type 3b [NiFe] hydrogenase. The locus contains the genes of the hydrogenase subunits hydADGB and of accessory proteins involved in enzyme maturation. Genes are drawn to scale. Locus tags (as found on MaGe) from left to right are as follows: AOA3F1_v2_1388, AOA3F1_v2_1387, AOA3F1_v2_1386, AOA3F1_v2_1385,AOA3F1_v2_1384, AOA3F1_v2_1383, AOA3F1_v2_1382, AOA3F1_v2_1381,AOA3F1_v2_1380.

**Fig S7.**
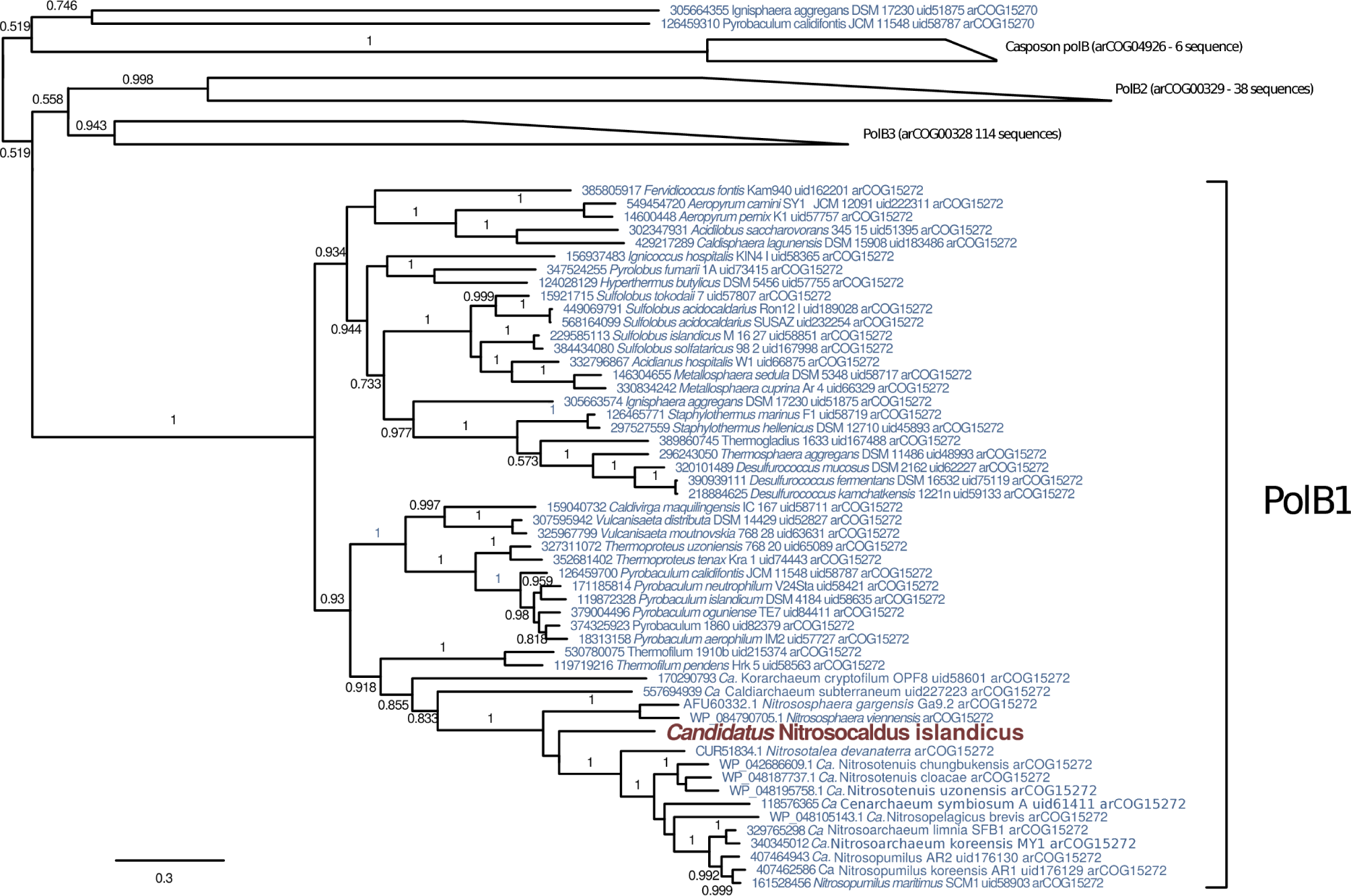
Unrooted approximate maximum likelihood tree showing placement of the PolB from *Ca.* Nitrosocaldus islandicus as a member of the PolB1 clade. The tree was calculated using FastTree^31^ on 213 sequences aligned with mafft^32^ (3058 aligned positions). Branch support greater than 0.5 is indicated on internal branches. PolB2, PolB3 and Casposon-related PolB (named according to 33) have been collapsed into right trapezoids in which the bases indicate shortest and longest terminal branches within each clade.

